# Neo-functionalization in *Saccharomyces cerevisiae*: A Novel Nrg1-Rtg3 chimeric transcriptional modulator is essential to maintain mitochondrial DNA integrity

**DOI:** 10.1101/2023.05.17.541047

**Authors:** Carlos Campero-Basaldua, James González, Janeth García-Rodriguez, Edgar Ramírez, Hugo Hernández, Beatriz Aguirre, Nayeli Torres-Ramírez, Dariel Márquez, Norma Sánchez, Nicolás Gómez-Hernández, Lina Riego-Ruiz, Claudio Scazzocchio, Alicia González

**Affiliations:** Departamento de Bioquímica y Biología Estructural, Instituto de Fisiología Celular, Universidad Nacional Autónoma de México, México; Laboratorio de Biología Molecular y Genómica, Departamento de Biología Celular, Facultad de Ciencias, Universidad Nacional Autónoma de México.; Departamento de Biología, Facultad de Química, UNAM, México City, Universidad Nacional Autónoma de México.; Laboratorio de Microscopía Electrónica Departamento de Biología Celular, Facultad de Ciencias, Universidad Nacional Autónoma de México.; Departamento de Genética y Biología Molecular, Instituto de Fisiología Celular Universidad Nacional Autónoma de México, México; División de Biología Molecular, Instituto Potosino de Investigación Científica y Tecnológica (IPICYT), México; Department of Life Sciences, Imperial College London, SW7 2AZ, UK; Université Paris-Saclay, CEA, CNRS, Institute for Integrative Biology of the Cell (I2BC), 91198, Gif-sur-Yvette, France

## Abstract

In *Saccharomyces cerevisiae*, the transcriptional repressor Nrg1 (Negative Regulator of Glucose-repressed genes) and the b/Zip transcription factor Rtg3 (ReTroGrade regulation) mediate glucose repression and mitochondria to nucleus signaling, respectively. Here we show a novel function for these two proteins, in which alanine promotes the formation of a chimeric Nrg1/Rtg3 regulator that represses the *ALT2* gene (encoding an alanine transaminase paralogue of unknown function) expression. A *NRG1/NRG2* paralogous pair, resulting from a post-wide genome, small scale duplication event, is extant in the *Saccharomyces* genus. Neo-functionalization of only one paralogue resulted in Nrg1, able to interact with Rtg3. Either *nrg1*Δ or *rtg3*Δ single mutant strains are unable to utilize ethanol and show a typical petite (small) phenotype on glucose. Neither of the WT genes complemented the petite phenotype, suggesting irreversible mitochondrial DNA damage in these mutants. Neither *nrg1*Δ nor *rtg3*Δ mutant strains express genes encoded by any of five polycistronic units transcribed from mitochondrial DNA in *S. cerevisiae*. This, and the direct measure of the mitochondrial DNA gene complement confirms that irreversible damage of the mitochondrial DNA occurred in both mutant strains and is consistent with an essential role of the chimeric Nrg1/Rtg3 regulator in mitochondrial DNA maintenance.

## Introduction

Gene duplication is a major evolutionary mechanism, which provides raw material for the generation of genes with novel or modified physiological roles (*Ohno 1970*; *Connant and Wolfe 2008*). Phylogenomic studies showed that in the ancestry of a clade of the Saccharomycotina, including *Saccharomyces cerevisiae* and other related yeasts (post-Whole Genome Duplication species, WGD), the occurrence of an interspecies hybridization event resulted in genome duplication (*Wolfe and Shields 1996; Marcet-Houben and Gabaldon 2015*). Selective retention and sub-functionalization of gene pairs derived from the hybrid ancestor has led to the functional divergence of the two paralogous copies (*DeLuna et al., 2001; Quezada et al., 2008; López et al., 2015; González et al., 2017*) or to neo-functionalization of one of the copies. The pairs of paralogous genes, that are extant in *S. cerevisiae*, could have both originated from one of the parental species (ohnologous pairs) or each one of the parental strains could have independently generated one member of the pair (homeologous genes). Duplication, however, could have arisen independently, preceding or following the WGD event. Duplication and neo-functionalization have been described for genes involved in primary metabolism, such as amino acid biosynthesis (*Escalera-Fanjul et al., 2017; Rojas-Ortega et al., 2018; González et al., 2017*) and also for genes encoding transcription factors (*Soussi-Boudekou et al., 1997; Poor et al., 2014; Merhej et al., 2015; Vick and Rine 2001*). *ALT1* and *ALT2* encode proteins with 65% identity, assumed to be paralogous alanine transaminases (*Chico et al., 1978*). Additional phylogenetic studies suggest that *ALT1* and *ALT2* originated in each one of the two parental strains which gave rise to the ancestral hybrid (*Huerta-Cepas et al., 2014*). *ALT1* encodes an alanine transaminase involved both in the main alanine biosynthetic pathway and in the sole alanine catabolic pathway. *ALT2*, despite its sequence similarity with *ALT1* (65% identity) is not an alanine transaminase, has no role in alanine metabolism; in fact, its physiological function is unknown (*Escalera-Fanjul et al., 2017; Rojas-Ortega et al., 2018*). Alanine plays a signaling role, as co-activator or co-repressor. Accordingly, *ALT1* expression is alanine induced (*García-Campusano et al., 2009*), while that of *ALT2* is alanine repressed. Sub-cellular localization has also diverged, since Alt1 is localized in the mitochondrial matrix while Alt2 is cytosolic (*Grandier-Vazeille et al., 2001*). Most interesting was the finding that Alt1 has been considered to be a moonlighting protein since besides its role in alanine catabolism, it contributes to the maintenance of mitochondrial DNA integrity and mitochondrial gene expression (*Márquez et al., 2021*).

Analysis of *ALT1* and *ALT2* gene promoters showed that both promoters carry an Nrg1 binding consensus sequence, while *ALT1* additionally includes an Rtg3 binding sequence. The present study shows, that although the *ALT2* gene promoter does not include an Rtg3 binding sequence, both Nrg1 and Rtg3 are necessary for alanine-dependent *ALT2* repression, which is alleviated in both *nrg1*Δ or *rtg3*Δ single mutants. These observations suggested the formation of a previously unreported Nrg1-Rtg3 chimeric complex. The roles of each of Nrg1 (*Vyas et al., 2005*) and Rtg3 (*Jazwinski et al., 2013*) as transcriptional regulators have been identified and thoroughly analyzed previously Nrg1 and its paralogue, Nrg2 regulate a group of stress responsive genes, functioning as transcriptional repressors (*Park et al., 1999; Vyas et al., 2003*). The first group of genes, which were found to be repressed by Nrg1/Nrg2 were some of those repressed by glucose. Further studies have shown that these two paralogues are also involved in responses to other environmental stresses such as: the alkaline pH stress response, hyperosmotic salinity response, negative regulation of pseudohyphal growth and biofilm formation (*Vyas et al., 2001*). The *NRG1*/*NRG2* pair arose as post-WGD small scale duplication event which occurred within the *Saccharomyces* genus, *NRG2* being the ancestral gene present in other Saccharomycotina (data in PhylomeDB 5 http://phylomedb.org/, (*Huerta-Cepas et al., 2011*)).

Rtg3 together with Rtg1 (not a paralogue of Rtg3) forms a complex which activates the retrograde signaling pathway from the mitochondria to the nucleus promoting the transcriptional activation of several genes (*Jazwinski et al., 2013*). The activation of the retrograde response extends the yeast replicative lifespan; it is enhanced and developed by the progressive decline in mitochondrial membrane potential during aging (*Liu and Butow 2006; da Cunha et al., 2015*). We here demonstrate that *ALT2* alanine-dependent repression necessitates the presence of the Nrg1-Rgt3, chimeric regulator, implying a novel physiological role of the Nrg1-Rtg3 chimera, non-extant in either Nrg1 or Rtg3 as has been found for Hap2-3-5-Gln3 and Gcn4-Gln3 hybrid chimeric regulators (*Hernández et al., 2011a; Hernández et al., 2011b*).

The possible existence of chimeric regulators was proposed by Forsburg and Guarente *(1989)*, when they established the presence of Hap4 as member of the HAP complex. They argued that, since the HAP complex was composed by a DNA binding domain involving three polypeptides (Hap2-3-5) and an independent activator sequence, included in the Hap4 protein, the combination of these peptides could generate novel chimeric regulators. The Hap2-3-5-Gln3 hybrid regulator, which allows Gln3 dependent *GDH1* transcriptional activation even in the presence of repressive nitrogen sources such as glutamine, constitutes a striking example (*Hernandez et al., 2011a*), which confirms the prediction of Forsburg and Guarente (*1989*). In all cells, multiple DNA binding proteins play crucial roles in the specificity of transcriptional responses; the existence of chimeric transcriptional modulators enlarges the repertoire of regulatory possibilities *(Forsburg and Guarente, 1989; Hernández et al., 2011a; Hernández et al., 2011b*).

This work presents inmunoprecipitation results demonstrating, the formation of an Nrg1/Rtg3 transcriptional complex, whose DNA binding domain is afforded by Nrg1, while both Rtg3 and Nrg1 are involved in chromatin organization of the *ALT2* promoter.

Our work on this chimeric regulator led to an unexpected result. The Nrg1/Rtg3 regulator has a far more crucial role than alanine mediated repression of *ALT2*. It is essential for the maintenance of mitochondrial DNA integrity. The absence of either component results in gross, irreversible, alteration of mitochondrial DNA. The formation of the Nrg1-Rgt3 complex implies a neo-functionalization of Nrg1, as the Nrg2 paralogue cannot function in the maintenance of mitochondrial DNA integrity. We argue that this neo-functionalization derives from the new acquired ability to form a chimeric complex with Rtg3.

## Results

### Nrg1/Rtg3 form a chimeric transcriptional complex whose assembly is promoted by alanine

As mentioned earlier, *ALT1* and *ALT2* arose from an interspecies hybridization which resulted in a WGD event (*Kellis et al., 2004; Marcet-Houben and Gabaldon 2015*). Alanine-dependent *ALT1* induction or *ALT2* repression is observed in the presence of alanine when cells are grown either on repressive nitrogen sources as glutamine or on non-repressive nitrogen sources such as proline or GABA (Figure 1A-C). The intracellular alanine pool is higher in GABA grown cells as compared to that found on either glutamine or proline (*Márquez et al., 2021*).

**FIGURE 1.**
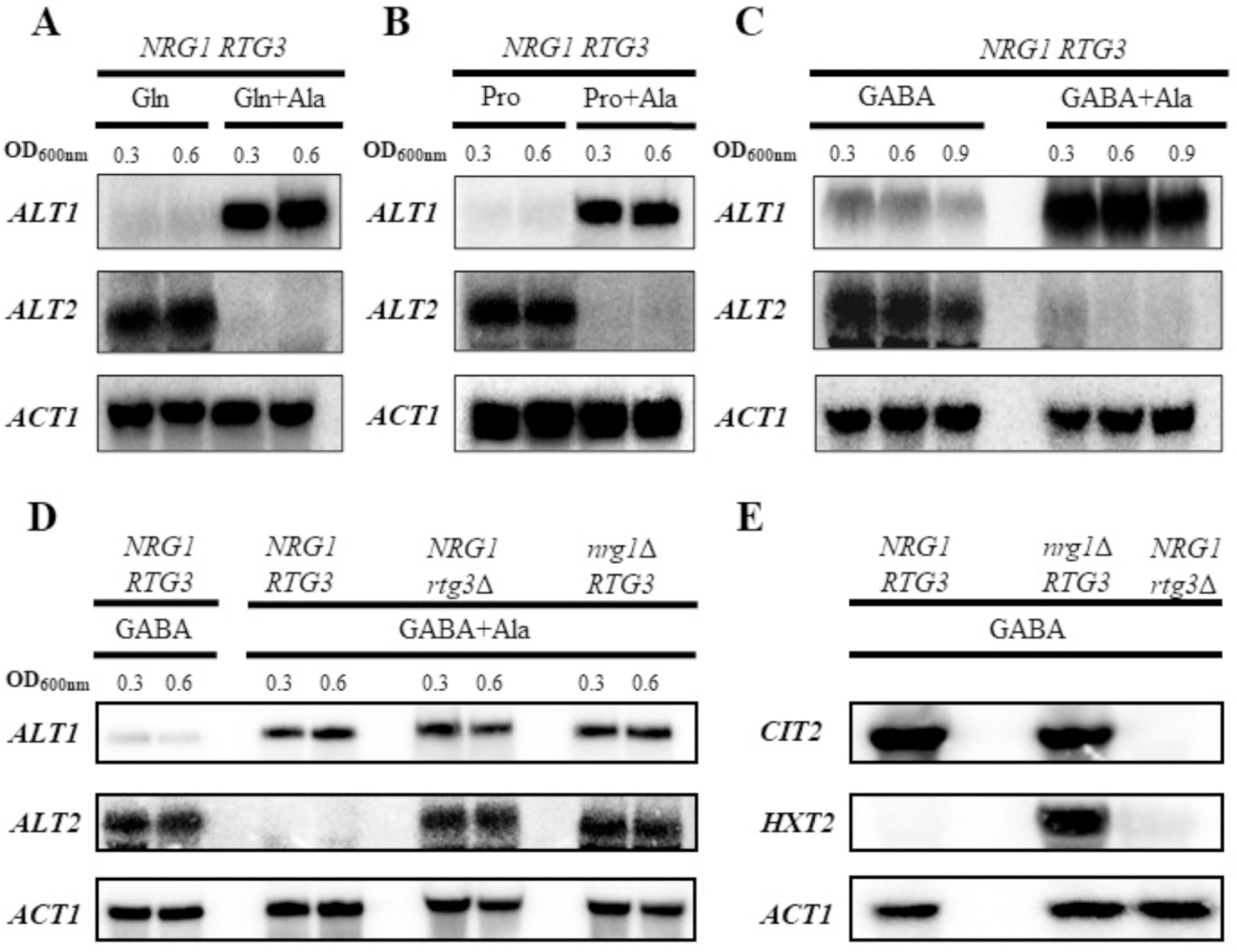
Alanine induces *ALT1* and represses *ALT2*. For Northern blot analysis, cultures were grown on glutamine (7 mM), proline (8 mM) and GABA (7 mM) with or without alanine (7 mM). (A) glutamine (Gln) and glutamine + alanine (Gln + Ala), (B) proline (Pro) and proline + alanine (Pro+Ala) (C) GABA (γ-aminobutiric acid) and GABA + alanine (GABA + Ala). When each culture reached an OD_600nm_ = 0.3 it was divided in two flasks and alanine was added to one of the cultures. Both cultures (with and without alanine) were incubated at 30°C for an additional 15 min. Then, samples of both cultures were taken for RNA extraction. Finally, a second sample of each culture was taken when cells reached an OD_600nm_ = 0.6 and RNA extraction was performed. (D) Cultures were prepared on GABA and GABA + Ala, samples were processed when cultures reached an OD_600nm_ = 0.3 and 0.6. (E) Cultures were grown on GABA to OD_600nm_ = 0.3 and expression of *CIT2* (citrate synthase) was monitored as a control of Rtg3 activity as a positive regulator, and *HXT2* (glucose transporter) as a target of Nrg1 as repressor. All samples were prepared for RNA extraction followed by Northern blot as described in Materials and Methods.

Analysis of the *ALT1* and *ALT2* gene promoters sequences showed that both promoters harbor DNA consensus binding sites for Nrg1, and that *ALT1* additionally bears a Rtg3 DNA consensus binding site. In order to analyze whether Nrg1 and/or Rtg3 could play a role in *ALT1* and *ALT2* expression, null *NRG1* and *RTG3* mutants were constructed. Northern analysis showed (Figure 1D) that both *nrg1*Δ or *rtg3*Δ mutants retained *ALT1* alanine-dependent induction, while *ALT2* repression did not occur in either mutant. We confirmed the previously described independent roles of *NRG1* and *RTG3* by monitoring the expression of *CIT2* (citrate synthase) and *HXT2* (high affinity glucose transporter) in each deletion mutant strain. As expected, *CIT2* expression is solely dependent on Rgt3 (*Jazwinski 2013*), while that of *HXT2* solely depends on Nrg1 (*Bae Lee et al., 2013*) (Figure 1E).

*ALT2* Nrg1 and Rtg3-dependent repression phenotype suggested that these regulators may form an Rtg3-Nrg1 chimeric repressor. In order to analyze this possibility, single and double *RTG3*-^TAP^ *NRG1*^-Myc13^ mutants were constructed as described in Materials and Methods. As Figure 2A shows, in a cell extract prepared from an *RTG3^-^*^TAP^ *NRG1*^-Myc13^ double-tagged strain immunoprecipitated with anti-Myc13 antibody, the presence of Rtg3^-TAP^ was revealed after Western analysis with anti-TAP antibody, indicating that Rtg3 and Nrg1 form part of the same complex. Co-immunoprecipitation was observed with either glucose or ethanol as carbon sources (Figure 2B). Interestingly, it was found that when cells were grown in the presence of alanine, co-immunoprecipitation enrichment was observed (Figure 2A and 2B). To further analyze this observation, experiments were carried out using increasing alanine concentrations, and as expected, alanine-dependent co-immunoprecipitation was proportionally higher in extracts obtained from cultures grown on 5 mM alanine as compared to those grown on 1 mM alanine (Figure 2C). These results suggest that alanine promotes the formation of the Nrg1-Rtg3 chimeric complex. However, Nrg1-^Myc13^ and Rtg3-^TAP^ co-immunoprecipitation is also observed in extracts prepared from cultures grown on GABA or proline in the absence of alanine (Figure 2C), suggesting that either the complex can be assembled without alanine or that the endogenously biosynthesized alanine pool can promote Nrg1-Rtg3 chimeric complex formation. Nrg1-^Myc13^ promoter occupancy was analyzed by chromatin immunoprecipitation (ChIP) experiments. Immunoprecipitates were obtained from Nrg1^-Myc13^ epitope-tagged strains with anti-Myc antibody. Amplification of *HXT2* and *GRS1* coding sequences were used as controls. As shown in Figure 2D, in the presence of alanine in the growth medium, Nrg1-^Myc13^ was recruited ten-fold more as compared to that observed in alanine absence to the *ALT2* promoter. This contrasts with result for the *HXT2* promoter in which Nrg1^-Myc13^ was similarly recruited in the presence of either 7 mM glutamine, proline, GABA or alanine. These results indicate that since higher amounts of Nrg1-Rtg3 complex are formed on alanine, a proportionally higher recruitment to the *ALT2* promoter must occur. However, even in the absence of exogenously added alanine the Nrg1-Rtg3 complex can be formed and recruited to the *ALT2* promoter.

**FIGURE 2.**
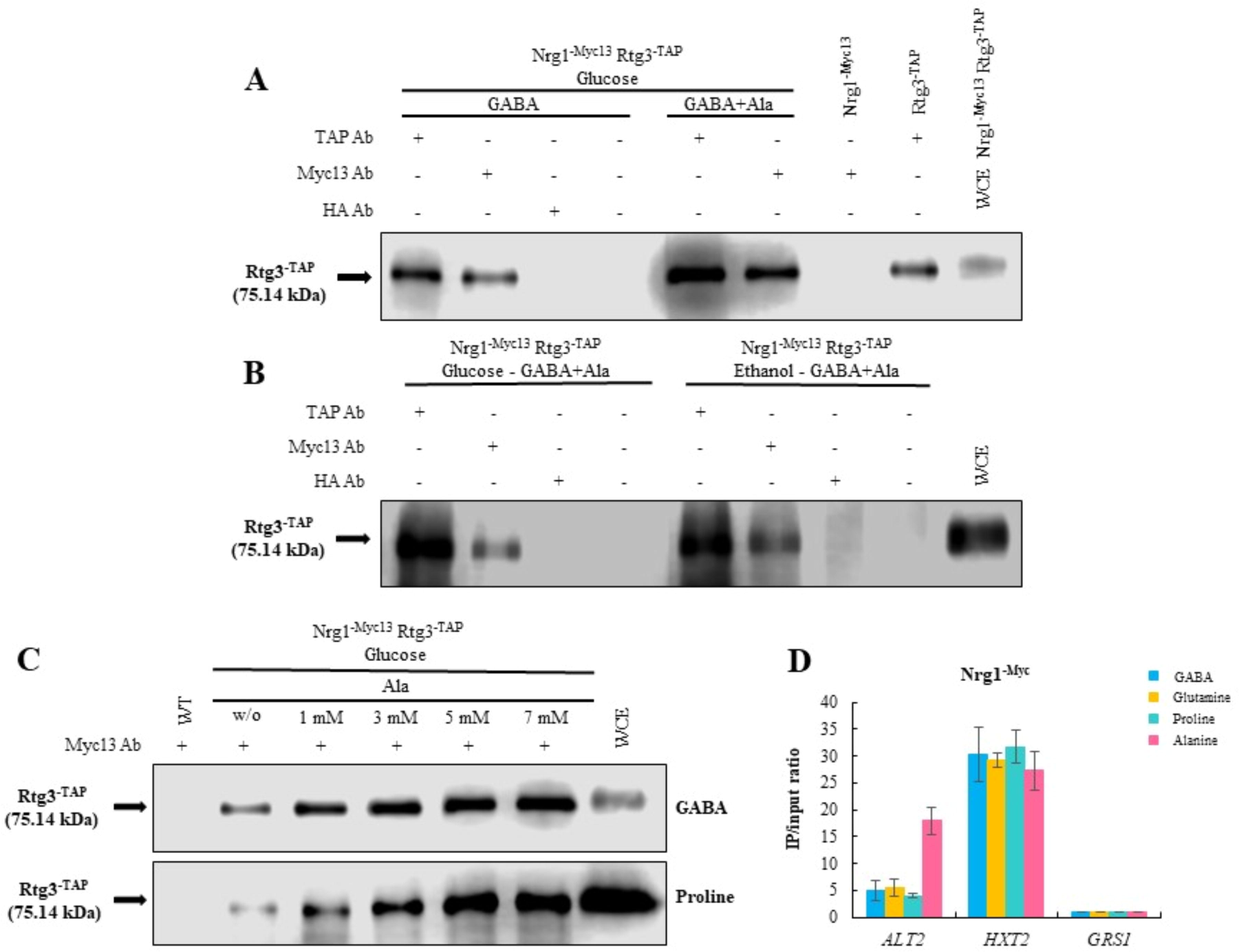
Rtg3 and Nrg1 form a transcriptional complex. Total protein samples were obtained from the double tagged Nrg1-^Myc-13^ Rtg3^-TAP^ strain, and single tagged Nrg1-^Myc-13^ or Rtg3^-TAP^ strains and processed as described in methods. Co-immunoprecipitation (CoIP): (A) compares the relative quantities of Rtg-^TAP^ coprecipitated in the presence or absence of alanine (7 mM). Extracts of the double mutant carrying gene fusions of the cognate genes with Myc-13 or TAP (see Materials and Methods) Nrg1-^Myc13^ Rtg3-^TAP^ were used for the immuno- and co-immunoprecipitation tests in the first six lanes. Single Nrg1-^Myc-13^ and Rtg3-^TAP^ tagged mutants grown on GABA (7 mM) + alanine (7 mM) were used as controls for Western blots revealed with anti-TAP. As a control, a Whole Cell Extract (WCE) of the double tagged strain grown on YPD was used to determine Rtg3-^TAP^ localization in the gel. Mouse hemagglutinin antibody (HA-Ab) was used as negative control, and the fourth lane did not carry antibodies. (B) Rtg3-^TAP^ and Nrg1-^Myc-13^ co-immunoprecipitation of extracts obtained from cultures grown on either MM 2% glucose or MM 2% ethanol. (C) Dependence of Rtg3 coprecipitated with Nrg1 on extracellular alanine concentration in extracts obtained from yeast cells grown on GABA (7 mM) or proline (8 mM). Immunoprecipitation tests were carried out separately with anti-Myc-13 (9E11 Santa Cruz Biotechnology), anti-TAP (Thermo Scientific) and anti-HA (F-7 Santa Cruz Biotechnology) as indicated. Samples (30 µl) were run on a 10% SDS-PAGE gel and Western blot analysis was carried out with anti-TAP antibody. A 6 µl sample of whole-cell extract (WCE) was used as a Rtg3-TAP control. (D) qChip assays were carried out on *ALT2*, *HXT2* and *GRS1* gene promoters in extracts obtained from the Nrg1-*^Myc-13^* tagged strain grown on MM in the presence of either 7mM GABA, glutamine, proline or alanine. *HXT2* (glucose transporter) was used as positive control and *GRS1* (glycyl-tRNA synthase) as negative control.

### DNA binding is dependent only on Nrg1, while both Nrg1 and Rtg3 activation domains contribute to chromatin remodeling

To analyze whether the Nrg1 DNA consensus binding domain played a role in Nrg1-Rtg3 binding to the *ALT2* promoter, chromatin immunoprecipitation experiments were carried out. Nrg1^-Myc13^ binding was increased three-fold in the promoter region of *ALT2* bearing the Nrg1 consensus binding sequence (TCCC). Rtg3^-Myc13^ binding is also increased four-fold in the same region, suggesting that the two activators are acting together in a complex, Nrg1 recruiting Rtg3 to the Nrg1 DNA binding domain. Consequently, chromatin immunoprecipitation was carried out in *nrg1*Δ *RTG3* and *NRG1 rtg3*Δ mutant strains. Figure 3B and 3C shows that Nrg1 binding is Rtg3 independent, while Rtg3 does not bind to the *ALT2* promoter in a *nrg1*Δ mutant background. Thus, Rgt3 promoter binding depends on Nrg1.

**FIGURE 3.**
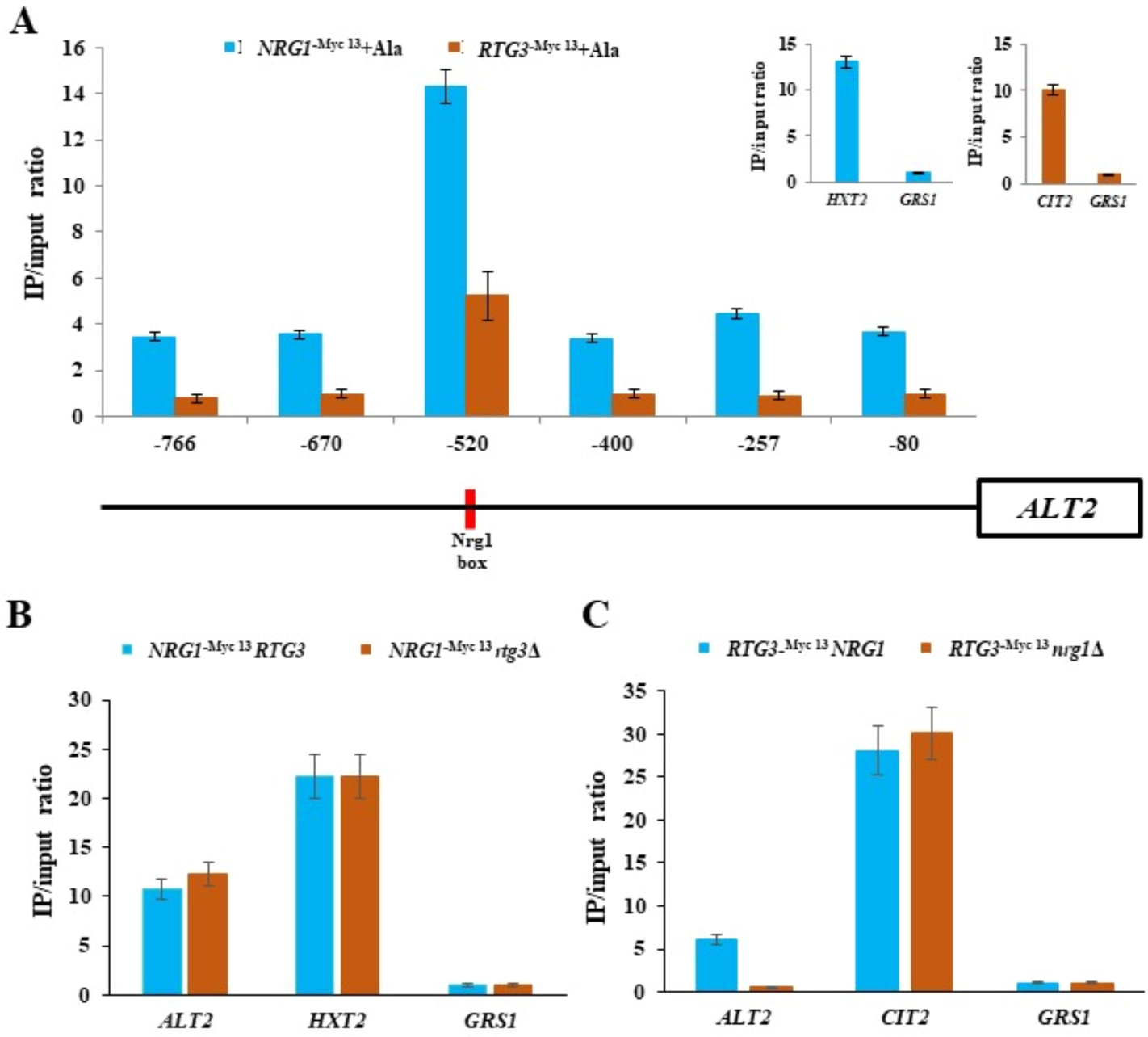
In the Nrg1-Rtg3 chimeric complex, the DNA binding domain is afforded by Nrg1. (A) qChip assays were carried out on the *ALT2* promoter region. The six DNA regions which were amplified after qChip are shown as vertical segments on the horizontal line representing various *ALT2* promoter regions. qChip assays were carried out with the anti-Myc-13 antibody (9E11, Santa Cruz Biotechnology) on WT strains containing Myc-13 epitope-tagged Nrg1-^Myc-13^ or Rtg3-^Myc-13^. Strains were grown on MM glucose (2%) + GABA (7 mM) and binding was analyzed by qChip as described in Materials and Methods. IP/input ratios were normalized with the *GRS1* promoter as negative control. *HTX2* (glucose transporter) and *CIT2* (citrate synthase) promoters were used as positive controls (shown in the right-side upper panels). In the *ALT2* depicted promoter, vertical red line shows Nrg1 consensus binding sequence. (B) qChip assays were carried out on the *NRG1-^Myc13^ RTG3* and *NRG1^-Myc-13^ rtg3*Δ. (C) qChip assay were carried out the *RTG3-^Myc-13^ NRG1* and *RTG3*-*^Myc-13^ nrg1*Δ. For (B) and (C) *HXT2* and *CIT2* promoters were respectively used as positive controls, IP/input ratios were normalized with the *GRS1* promoter as negative control. Data are presented as the average of three independent experiments.

Analysis of *ALT2* chromatin organization showed the *ALT2* promoter is condensed when the wild type strain is grown on GABA as sole nitrogen source (Figure 4A). When GABA and alanine were simultaneously provided the *ALT2* promoter organization is further compacted (Figure 4A), in line with the repressive role of alanine. Conversely, the *ALT1* promoter shows a more relaxed organization in the presence of alanine, in accordance with the co-activator role of alanine in *ALT1* expression (*Márquez et al., 2021*). In both, *nrg1*Δ or single *rtg3*Δ mutants, *ALT2* promoter chromatin organization was relaxed, in accordance to the *ALT2* de-repressed expression seen in these strains (Figure 4B and 4C, and Figure 1D).

**FIGURE 4.**
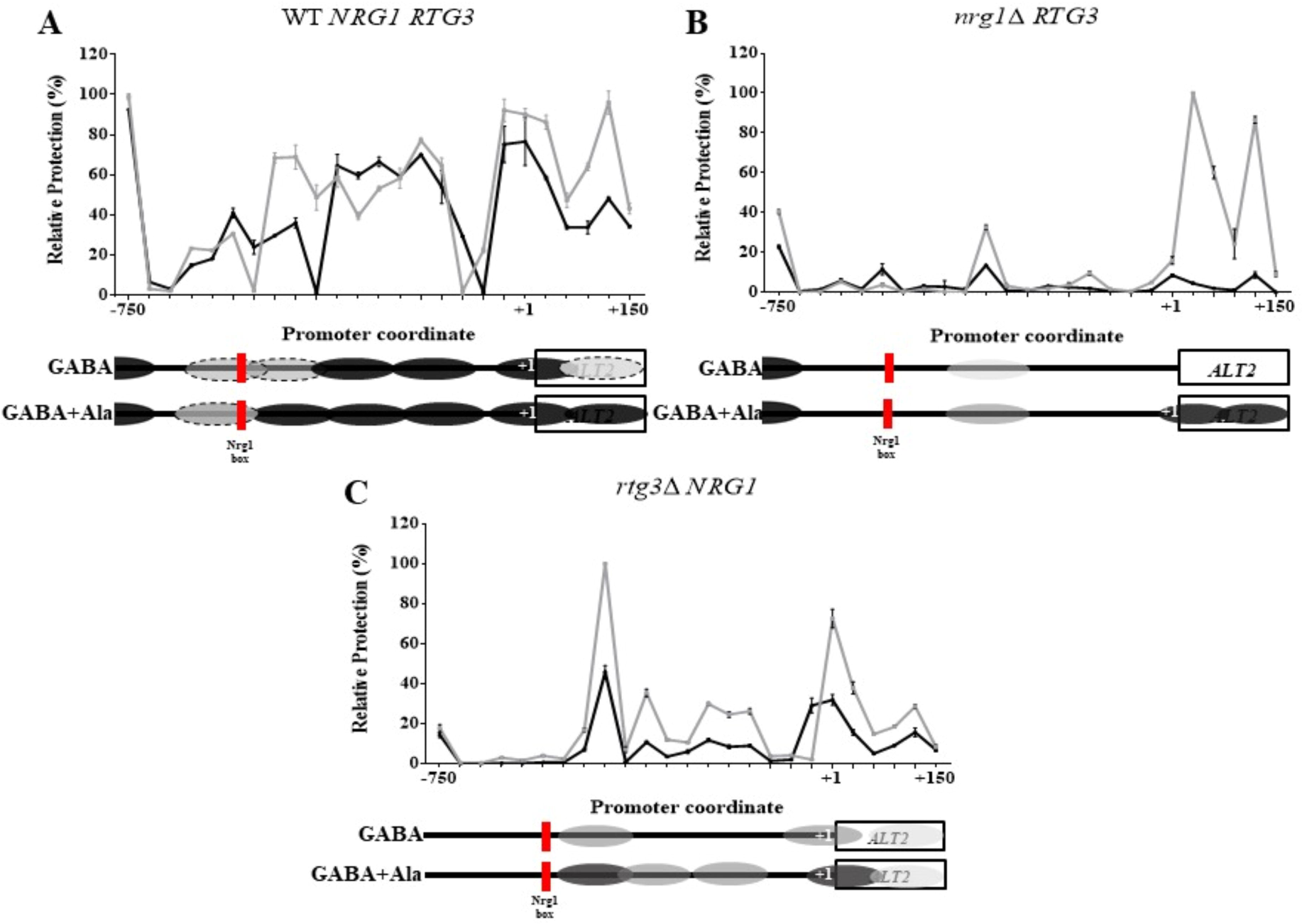
Both, Nrg1 and Rtg3 are necessary for chromatin rearrangement in the *ALT2* promoter. NuSA (Nucleosome Scanning Assay) samples were obtained as follows, wild type (WT) or mutant strains were grown on MM glucose (2%) + GABA (7 mM) or glucose (2%) GABA (7 mM) + alanine (7 mM). When each culture reached an OD_600nm_ = 0.6 samples of both cultures were taken for nucleosome scanning assay, as described in Materials and Methods. Mean values of three independent experiments are shown. Black lines show chromatin organization observed on GABA and gray lines on GABA + ala grown cells. In the *ALT2* depicted promoter, vertical red line shows Nrg1 consensus binding sequence, black ovals over *x-*axis indicate firmly positioned nucleosomes and dotted ovals depict fuzzy nucleosomes. (A) WT, (B) *nrg1*Δ and, (C) *rtg3*Δ.

### Absence of either Nrg1 or Rtg3 results in complete loss of mitochondrial DNA and respiratory metabolism

It has been previously reported that *rtg3*Δ mutants show glutamate braditrophy, since Rtg3 absence results in loss of the retrograde response, hindering the transcriptional activation of *CIT1*, *ACO1* and IDH1/2, which allow glutamate biosynthesis (*Jazwinski 2013*). Figure 5A and 5B shows that, as expected, *rtg3*Δ mutants are glutamate braditrophs which when transformed with pRS416−*RTG3* strains recover glutamate prototrophy, confirming the recovery of the *RTG3* wild type phenotype.

**FIGURE 5.**
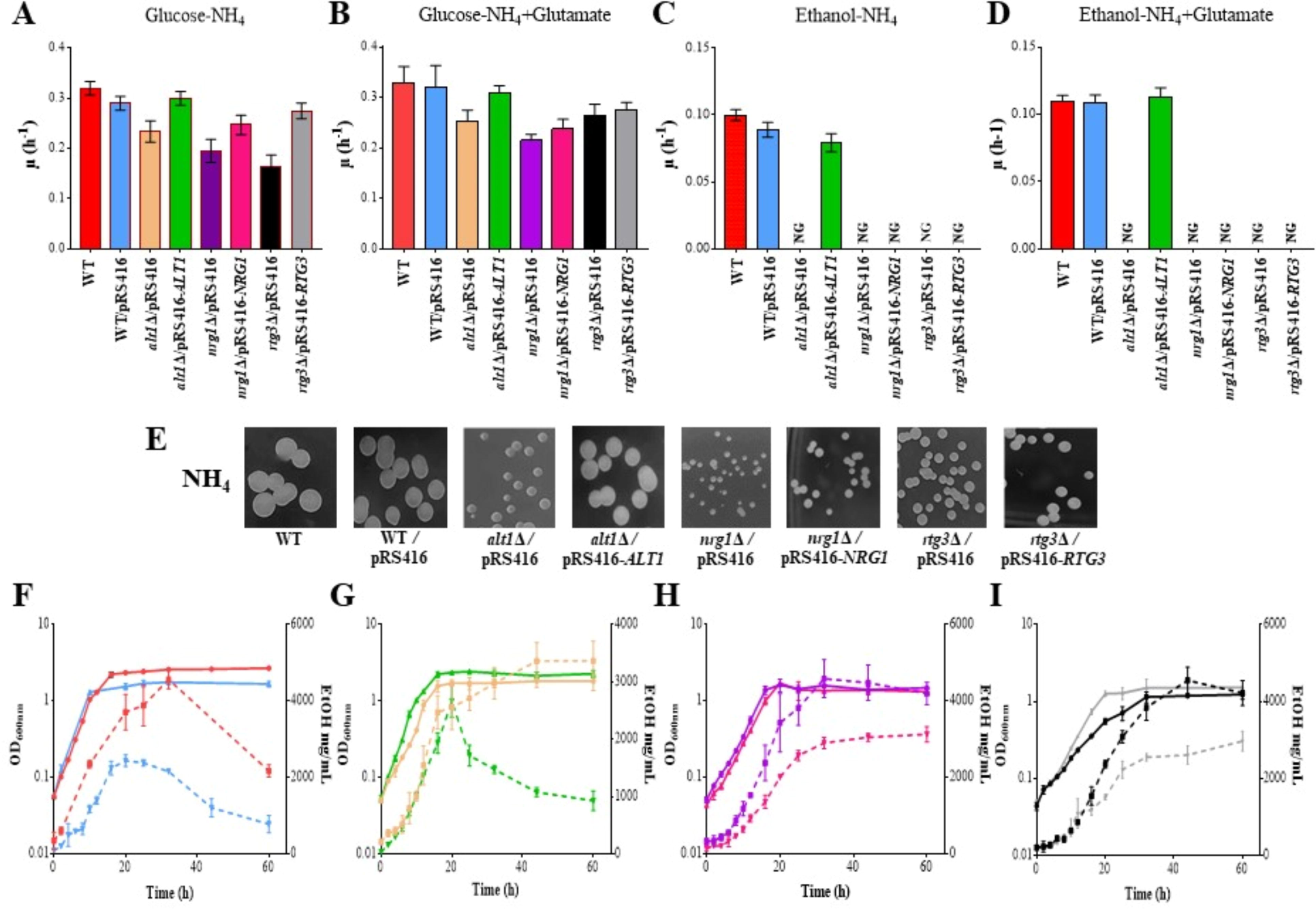
*nrg1*Δ and *rtg3*Δ mutants show a petite phenotype even when transformed with their cognate *NRG1* or *RTG3* wild type genes. (A) Specific growth rate on MM glucose (2%)-ammonium (40 mM), (B) specific growth rate on MM glucose (2%)-ammonium (40 mM) + glutamate (5 mM); (C) specific growth rate in MM ethanol (2%)-ammonium (40 mM); (D) specific growth rate on ethanol-ammonium + glutamate (5 mM). Results of three independent experiments are presented. (E) Colony size in glucose + ammonium. Plates were incubated at 30°C during 3 days on glucose ammonium (NH_4_). (F-M) Cells were cultured to monitor biomass (solid lines) and ethanol concentration (dotted lines) on MM glucose + ammonium. Each line represents the average of three independent experiments.

The presence of the transforming plasmid was monitored by the complementation of the uracil auxotrophy present in the receiving mutant strains.

Neither Rtg3 nor Nrg1 transformants recovered the capacity to grow on ethanol as carbon source with or without glutamate as nitrogen source (Figure 5C and 5D), suggesting that mitochondrial function has been irreversibly impaired.

As expected, *nrg1*Δ and *rtg3*Δ mutants showed a petite phenotype, forming small colonies when grown on glucose as carbon source, and this phenotype was not complemented in *nrg1*Δ/pRS416-*NRG1* or *rtg3*Δ/pRS416-*RTG3* transformed strains confirming that absence of Nrg1 or Rtg3 has resulted in an irreversible damage to the mitochondrial function (Figure 5E).

Analysis of the ability to generate and consume ethanol by the *nrg1*Δ and *rtg3*Δ mutants was determined, showing that *nrg1*Δ and *rtg3*Δ mutant strains could produce ethanol but were unable to consume it. Furthermore, the transformed strains *rtg3*Δ/pRS416-*RTG3* and *nrg1*Δ/pRS416-*NRG1* did not recover the capacity to consume ethanol, (Figure 5F-M). As a control, ethanol production and consumption were analyzed in the petite strain *alt1*Δ (*Márquez et al., 2021*) and in the cognate strain transformed with pRS416-*ALT1*. In this case, *alt1*Δ harbouring the empty plasmids pRS416 produced but did not consume ethanol, while a strain transformed with pRS416-*ALT1* recovered the wild type capacity to produce and consume ethanol and does not show a petite phenotype. To further analyze Rtg3 and Nrg1 roles on the regulation of mitochondrially encoded genes, we determined expression of five mitochondrial DNA-encoded genes. The *S. cerevisiae* mitochondrial DNA encodes cytochrome oxidase subunits I, II y III (*COX1, COX2, COX·3*), apocytochrome b (*COB1*) and three ATP synthase subunits (*ATP6, ATP8, ATP9*) (*Turk et al., 2013*). Since mitochondrial genes are expressed as polycistronic transcripts, we determined the expression of five of the above mentioned genes (*COX2, COX3, COB1*, *ATP6* and *ATP9*) representing different transcriptional units, since *COX1*, *ATP8* and *ATP6* belong to the same polycistronic unit, determining *ATP6* expression informs on the expression of this whole polycistronic unit (*Turk et al., 1999*). Thus, we analyzed the five polycistronic units encoding proteins from the mitochondrial genome. As shown in Figure 6, *COX2* and *COX3*, both forming part of respiratory complex IV (*Geier et al., 1995*); *COB1* which constitutes a subunit of the ubiquinol-cytochrome c reductase complex (*Hunte et al., 2000*); *ATP6* which encodes the subunit a of the F0 sector of FIFO ATP synthase (*Turk et al., 1999*) and *ATP9* encoding the F0-ATP synthase subunit (*Payne et al., 1993*) expression was observed in the wild type strain and completely abolished in either one of *rtg3*Δ or *nrg1*Δ mutants grown on glucose-GABA or glucose-GABA + alanine. We further analyzed the expression of the nuclearly-encoded *COX8* and *COX6* genes, which constitute the VI and VIII subunits of cytochrome c oxidase, which forms part of Complex IV, the terminal member of the mitochondrial inner membrane electron transport chain (*Geiger et al., 1995*) and that of *ALT2* encoding the *ALT1* paralogous of unknown function. As Figure 6 shows, the expression of *ALT2* is repressed in the wild type strain and de-repressed in *nrg1*Δ and *rtg3*Δ strains. In cultures of strains grown on either GABA or GABA + alanine. *COX6* and *COX8* expression is repressed in alanine but it is not regulated by Nrg1 or Rtg3

**FIGURE 6.**
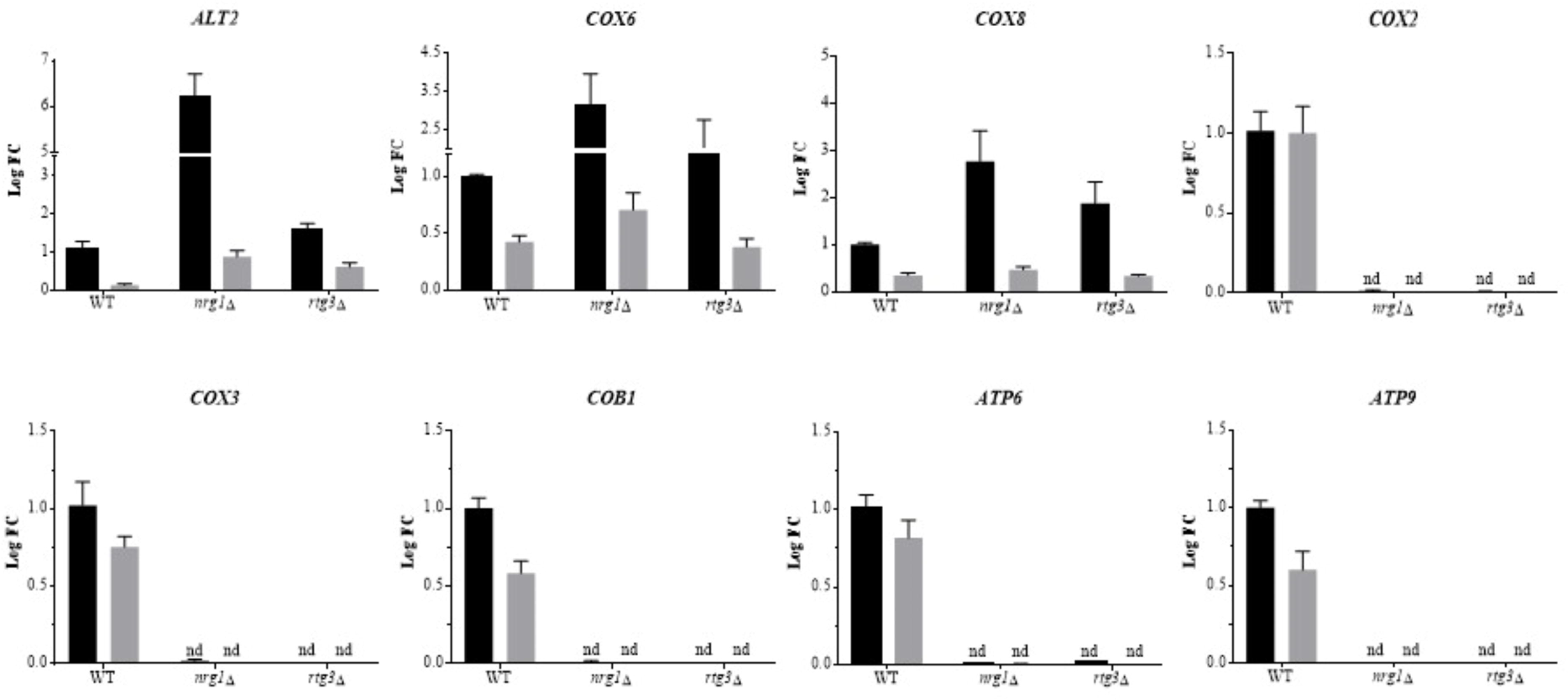
Nrg1-Rtg3 complex is necessary for mitochondrial DNA expression. Strains were grown on MM glucose (2%) - GABA (7 mM) or MM glucose (2%) – GABA (7 mM) + alanine (7 mM) as a nitrogen source and collected at OD_600nm_ = 0.3. Gene expression was measured by qPCR using 18S as constitutive control by the 2^-ΔΔCT^ method. qPCR analysis shows *ALT2, COX8* and *COX6* nuclear genes expression in the *NRG1 RTG3*, *nrg1*Δ *RTG3* and *NRG1 rtg3*Δ strains, grown on GABA (7 mM) or GABA (7mM) + alanine (7mM), indicating these are repressed by Nrg1 and Rtg3. The *COX2*, *COX3*, *COB1*, *ATP9* and *ATP6* mitochondrial genes expression is absent in *rtg3*Δ and *nrg1*Δ mutants. Black bars, MM glucose – GABA; gray bars, MM glucose – GABA + alanine. nd, not detected.

We then determined the ratio of mtDNA/nDNA using *COX6* as a nuclear control. As shown in Figure 7 A and 7B, when DNA was quantified in samples obtained from glucose-GABA or glucose-ammonium grown cells, the mtDNA/nDNA ratio showed that mtDNA was completely absent even in the strains transformed with the cognate WT genes.

**FIGURE 7.**
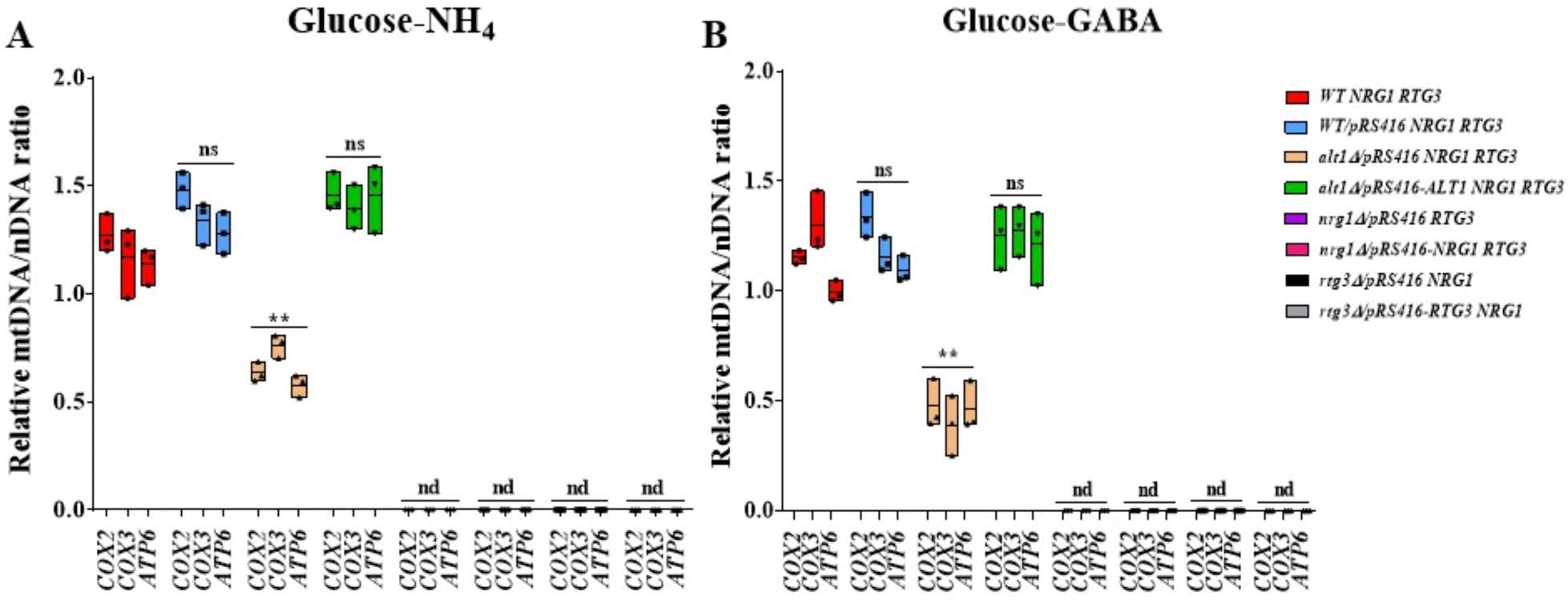
Both *nrg1*Δ and *rtg3*Δ result in drastic depletion of mitochondrial DNA. Cells were harvested on (A) MM glucose (2%) + ammonium (NH_4_, 40 mM) or (B) MM glucose (2%) + GABA (7 mM) and collected at OD_600nm_ = 0.6. Estimation of mtDNA/nDNA ratio by qPCR using *COX2, COX3* and *ATP6* as mitochondrial genes; this determination is relative to the *COX6* encoded nuclear gene. Boxes show individual values (points) and the average (middle line) + SD of three independent experiments, Statistical analysis were done using Anova two-way. Asterisks indicate a significant difference **= P < 0.005. ns, non-significant; nd, not detected.

Thus, both Nrg1 and Rgt3, acting in a complex, play a crucial role in maintaining mitochondrial DNA integrity and/or stability. Consequently, we looked at the appearance of mitochondria as monitored by MItotracker green. Figure 8 shows that the appearance of mitochondria on both, the wild type and *alt1*Δ strain complemented with *ALT1* show a typical filamentous organization, while *alt1*Δ shows a clearly diminished filamentous organization (*Márquez* et al., 2021). This contrasts with the fragmented pattern observed in both, *nrg1*Δ and *rgt3*Δ strains, whether complemented or not with their cognate WT genes (Figure 8), confirming that the absence of either member of the chimeric regulator results in irreversible damage to mitochondria, in line with loss of mitochondrial DNA.

**FIGURE 8.**
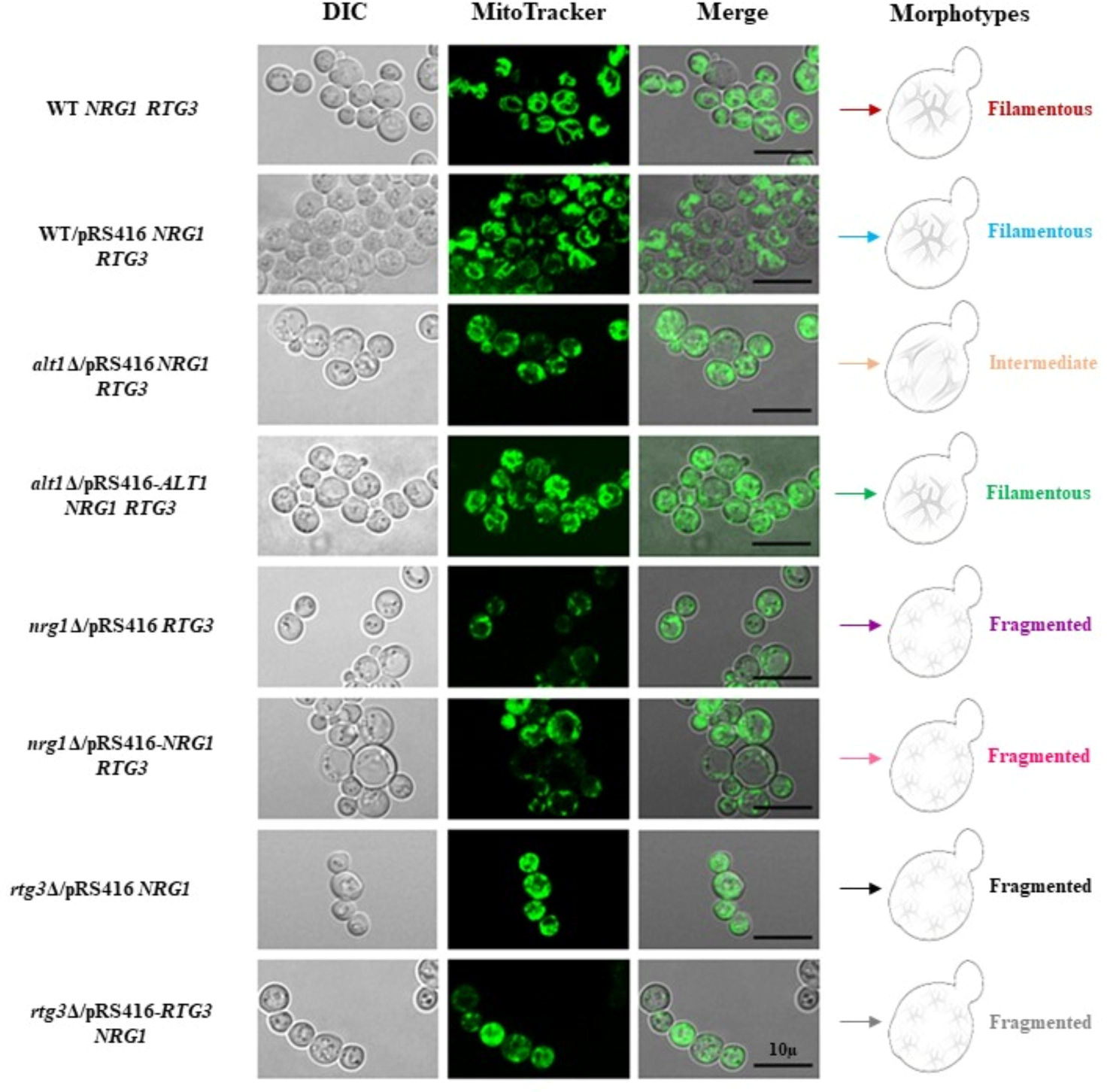
*nrg1*Δ and *rtg3*Δ show a drastically altered mitochondrial morphology. Mitochondrial staining with MitoTracker Green FM 9074 (Molecular Probes) to determine the morphology of mitochondria seen in each strain. Black line scale corresponds to 10 μ. Morphotypes show schematic representation of mitochondrial organization in the different strains.

### Modelling of the Nrg1/Rtg3 complex, the unicity of *NRG1*

We attempted to investigate the structure of the Nrg1/Rtg3 complex through its modelling using alpha-fold (see Materials and Methods). Figure 9 shows the most probable complex, which was deduced. Interestingly, neither of the components of the complex can be superimposed to the alpha-fold models (available in the data base *NRG1*AF-A0A815XE22-F1-model_v4.pdb and *RTG3* AF-A0A815WEC9-FI-model_ v4.pdb) through the Multiseq plugin available in VMD (*Humphrey et al., 1996*), which suggests that the formation of the complex alters substantially the tridimensional configuration of both components (Figure 9). We analyzed the possible contacts between the components of the complex (*Varadi et al., 2022*). These lie in two patches. A number of contacts are extant between two intrinsically disordered stretches (please see Figure 9 legend). An additional contact is made by Lys327 of Rtg3 with Asp164 and Glu165 of Nrg1 (Supplementary Table 1). These contacts provide a clear rationale for the unique specificity of Nrg1. *NRG1* and *NRG2*; show a global 50.46 identity; however, our results show clearly that *NRG2* cannot substitute *NRG1* as a component of the chimeric regulator. Nrg1 neo-functionalization resulted in the acquisition on the pertinent amino acid residues supporting Nrg1-Rtg3 interactions. Nrg1 and Nrg2 differ between residues 142 (Met in Nrg1) and 171 (Arg in Nrg1) where the main patch of contact residues is located. Moreover, the Asp164 and Glu165 which form a salt bridge and H bonds with Lys327 of Rtg3 are absent in Nrg2. The predicted alpha-fold models of Nrg1 and Nrg2 are also substantially different (supplementary Figures 1 and 2).

**FIGURE 9.**
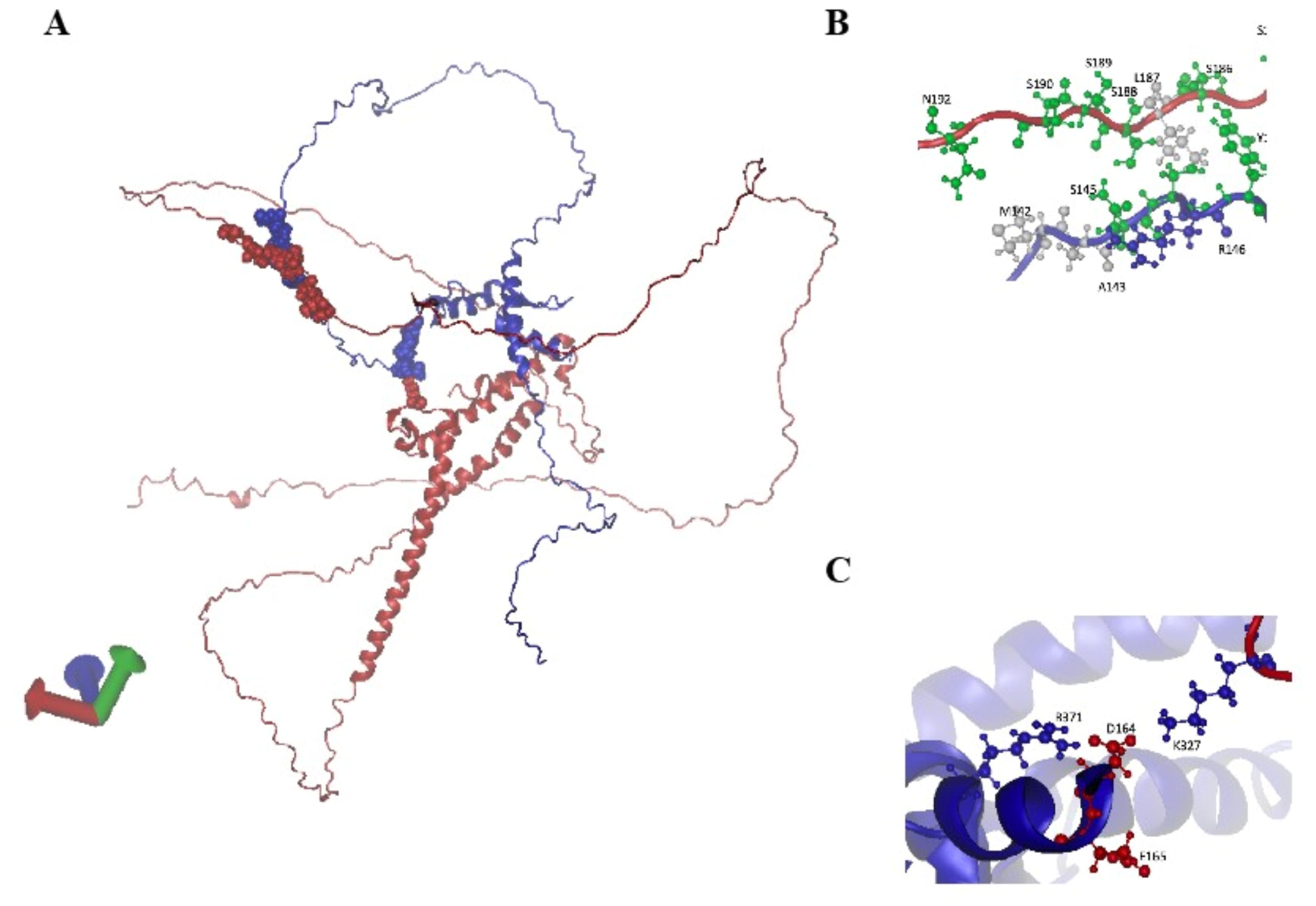
Modelling the Nrg1/Rtg3 interaction. To the left, Panel A most likely model of the Nrg1/Rtg3 hybrid molecule obtained by ColabFold v1.5.2: AlphaFold2 using MMseq (https://colab.research.google.com/github/sokrypton/ColabFold/blob/main/AlphaFold 2.ipynb#scrollTo=ADDuaolKmj) (*Mirdita et al., 2022*). Blue chain, Nrg1; Red Chain, Rtg3. The residues in the two contact patches (see below) are shown for clarity as VDW (space filling). To the right, Panels B and C, detailed residues involved in the two contact patches as determined with the MAPIYA contact server (https://mapiya.lcbio.pl/) (*Badaczewska-Dawid et al., 2022*).

**FIGURE 10.**
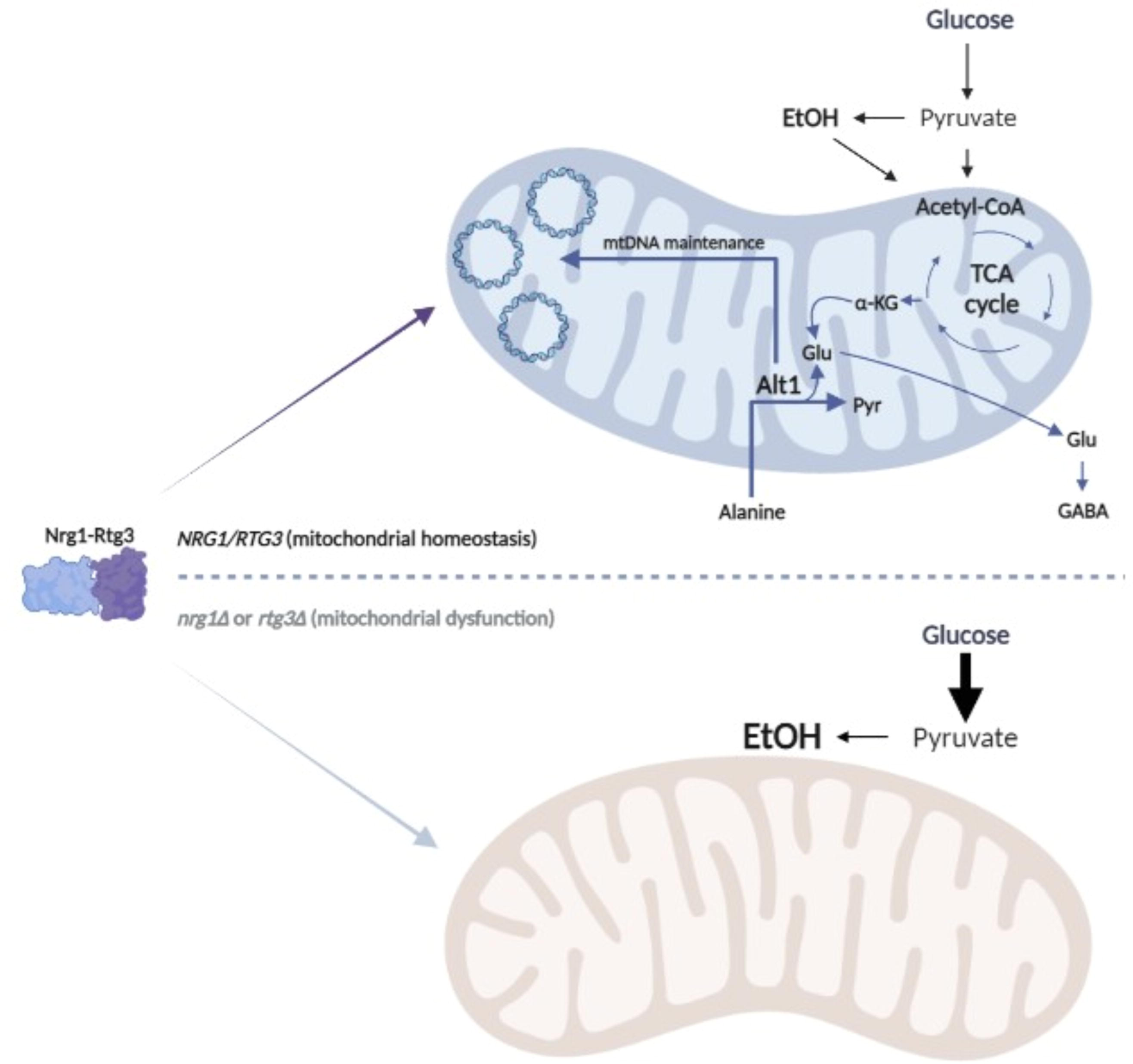
Nrg1-Rtg3 chimeric transcriptional modulator maintains mitochondrial homeostasis in *S. cerevisiae*. After neofunctionalization, Nrg1 gained amino acid residues which allow assembly of the Nrg1-Rtg3 novel chimeric modulator, which maintains mitochondrial homeostasis due to a yet unknown mechanism. Alanine catabolism through the Alt1 moonlight protein (alanine transaminase/mtDNA maintenance) provides glutamate (Glu), which feeds the GABA shunt pathway. Our results indicate that in the absence of Nrg1 (*nrg1*Δ) or Rtg3 (*rtg3*Δ), the chimeric Nrg1-Rtg3 is not assembled causing mitochondrial disfunction which results in lack of mitochondrial DNA (mtDNA), null respiratory metabolism, and fragmented mitochondria (bottom panel), which affects essential mitochondrial functions such as stress response, apoptosis, mitophagy or aging. Ethanol (EtOH), α-ketoglutarate (α -KG), glutamate (Glu), and pyruvic (Pyr). Image created with BioRender.com.

### Phenotypic Characterization of *nrg2*Δ mutants

The clear phenotype of *nrg1*Δ mutants implies that *NRG2* cannot fulfill the *NRG1* physiological function. We did, however, an *nrg2*Δ single mutant and an *nrg1*Δ *nrg2*Δ double mutant, as described in Materials and Methods. As expected, an *nrg2*Δ mutant grew on both ethanol or glucose as carbon sources showing wild type phenotype, while the double mutant *nrg1*Δ *nrg2*Δ showed a petite phenotype on glucose and did not grow on ethanol as sole carbon source (Supplementary Figure 3).

## DISCUSSION

### Alanine as a metabolic signal promoting organization of the Nrg1-Rtg3 modulator

Alanine aminotransferases catalyze the reversible transamination between alanine and U-ketoglutarate leading to pyruvate and glutamate. At least two alanine aminotransferase isozymes generated through independent gene duplication events, occur frequently in animals, plants, yeasts and bacteria, and they are proposed to be involved in the main pathways of both alanine biosynthesis and catabolism (*Gatehouse et al., 1967; Chico et al., 1978; Wang et al., 1987; Umemura et al., 1994; De sousa and Sodek 2003; Liu et al., 2008*). Amino acid metabolism has been thoroughly studied in *S. cerevisiae*, however, no alanine auxotrophs have ever been isolated in this yeast. Moreover, although Alt1 is the main alanine biosynthetic enzyme contributing to 80% of the alanine intracellular pool, *alt1*Δ mutation does not lead to alanine auxotrophy, indicating that at least one additional pathway for alanine biosynthesis must be extant in this yeast (*García Campusano et al 2009; Peñalosa-Ruiz et al., 2012*). There are two candidates for an alternative alanine biosynthesis pathway; the glutamine-pyruvate aminotransferase (*Cooper and Meister 1977; Calderon et al., 1985; Soberón and González, 1987; Soberón et al., 1989; Duran et al., 1995*), and the Uga1 encoded GABA transaminase, where the resulting succinate semialdehyde is readily oxidized through Uga2 rendering alanine biosynthesis irreversible (*Márquez et al., 2021*). It is thus likely that in *S. cerevisiae* an alanine intracellular pool is permanently available (*Márquez et al., 2021*), which could facilitate assembly of the chimeric Nrg1-Rtg3 regulator we have described here.

Alanine plays a role as an *ALT1* positive regulator inducer and an *ALT2* co-repressor while it also represses the expression of the genes involved in the GABA shunt (*Márquez et al., 2021*). As the formation of The Nrg1-Rtg3 complex is enhanced in the presence of alanine, this suggests that this amino acid could play an additional regulatory role in facilitating and/or promoting the assembly of this chimeric regulator. Our results show that even on proline as sole nitrogen source, a condition in which the alanine intracellular pool is reduced (*Márquez et al., 2021*), the Nrg1-Rtg3 chimeric complex is formed (Figure 2C). Additionally, we show that *ALT2* Nrg1-Rtg3 dependent repression is observed even on cultures grown on GABA in the absence of added alanine (Figure 6), further supporting that Nrg1-Rtg3 assembly could be mediated by the intracellular alanine pool.

### Nrg1-Rtg3-dependent regulation is essential to maintain DNA integrity and mitochondrial function

We have demonstrated that Nrg1 and Rtg3 form a chimeric transcriptional regulator. Analysis of the homologous pair *NRG1* and *NRG2*; show a global 50.46 identity. However, *NRG2* cannot substitute the *NRG1* encoded component of the chimeric regulator as *nrg2*Δ mutants do not share *nrg1*Δ or *rtg3*Δ mutant phenotypes. Indeed, the neofunctionalized Nrg1 acquired the amino acid residues that can mediate Nrg1-Rtg3 interaction, and which are absent in Nrg2. When the Nrg1-Rtg3 chimera cannot be assembled, as in the *nrg1*Δ or *rtg3*Δ mutants, mitochondrial DNA is irreversibly damaged, resulting in extreme mitochondrial fragmentation (Figure 8), thus cells cannot carry out respiratory metabolism. When these mutants are transformed with the *NRG1* or *RTG3* wild type genes the capacity to carry out respiratory metabolism is not recovered, since mitochondrial DNA has been lost. The fact that either a *nrg1*Δ or a *rtg3*Δ mutant results in loss of mitochondrial DNA strongly suggests that this phenotype results from the absence of the Nrg1-Rtg3 chimeric complex. This is a hitherto undescribed function, not carried out by the individual, un-complexed, Nrg1 or Rtg3 regulators. It is unlikely that his new role of the Nrg1-Rtg3 chimeric complex in maintaining mitochondrial DNA integrity could be related to its role in repressing *Alt2*, considering that *alt2*Δ mutants do not show any obvious phenotype. We favor the idea that Nrg1-Rtg3 complex plays additional regulatory roles, positive or negative, on a gene or genes crucially involved in mitochondrial DNA replication and/or maintenance.

Formation of chimeric complexes such as Hap2-3-5-Gln3 (*Hernandez et al., 2011a*) and the herein described Nrg1-Rtg3, supports the notion that extant elements can be recruited to novel functions thorough their interactions (*Baldwin and Krebs, 1981)*.

Besides the traditional mitochondrial role in energy transformation, this organelle is involved in many other crucial functions. Mitochondria are vital in sustaining cellular and organismal pathways that determine the direction metabolism can follow, develop pertinent stress responses, and cellular fate (*Shen et al., 2022*). To fully accomplish these tasks, networks of intracellular communication are extant, which depend on diverse and numerous molecular cascades. Indeed, feedback from mitochondria is required to coordinate mitochondrial biogenesis and/or removal by mitophagy during the division cycle (*Knorre et al., 2016*) and a coordinated regulation of these processes by hybrid transcriptions factors such as the Nrg1-Rtg3 complex can allow cellular adaptations to stress and other environmental modifications (*Knorre et al., 2016; Shen et al., 2022; Balaban et al., 2005*).

When glucose becomes limiting, *S. cerevisiae* cells undergo a diauxic shift, turning to respiratory metabolism. Under this condition, *S. cerevisiae* cells rely on aerobic energy production to support growth. Absence of either component of the chimeric Nrg1-Rtg3 regulator results in loss of respiratory metabolism leading to a fully fermentative metabolism. Such facultative property of *S. cerevisiae* allowed us to uncover this novel system involved in the maintenance of mitochondrial DNA integrity It would be interesting to see if this same system is present in obligatory aerobic yeasts and the evolution of alternative systems in other eukaryotes.

## CONCLUSION

Metabolic adaptation to diverse and new environments results in complex modifications of the physiology of cells, allowing them to turn on relevant response pathways. Although all organisms include a vast array of DNA binding proteins participating in adaptive transcriptional responses, the formation of chimeric modulators, combining different DNA binding and activation domains, can expand the repertoire of activators or repressors allowing the development of novel and more complex responses (*Hernández et al., 2011a; Hernández et al., 2011b*). This is the case of the Nrg1-Rtg3 chimeric modulator we have described here as essential to maintain respiratory metabolism, and mitochondrial-dependent energy production.

## Materials And Methods

### Strains

Tables 1 and S1 describe the characteristics of the strains and plasmids used in the present work. CLA1 *ura3 leu2* construction has been previously described (*Valenzuela et al., 1998*). CLA1-2 (*ura3* leu2::LEU2) (*Quezada et al., 2008*) strain was transformed with plasmid pRS416 obtaining *CLA1-2-A/pRS416* strain used as a control harboring the empty plasmid. Construction of strain CLA11-713 (*nrg1*Δ*::kanMX4 ura3 leu2::LEU2*) has been previously described (*González et al., 2017*). The CLA11-714 (*nrg2*Δ::*kanMX4 ura3*Δ *leu2*Δ) was obtained by gene replacement. A PCR-generated *kanMX4* module was prepared from plasmid pFA6a (Table S1) using R5 and R6 deoxyoligonucleotides (Table S2a). The CLA11-715 double mutant (*nrg1*Δ::*natMX4*; *nrg2*Δ:: *kanMX4 ura3*Δ *leu2*Δ) was constructed as follows. The *kanMX4* module from the (*nrg1*Δ*::kanMX4 NRG2 ura3 leu2*) was replaced by the *natMX4* cassette which confers resistance to the nourseothricin antibiotic (*Goldstein and McCusker 1999*). The *natMX4* module used for transformation was obtained from plasmid p4339 (Table S1) using R3 and R4 deoxyoligonucleotides (Table S2a). The module obtained of *nrg1*Δ::*natMX4* was then transformed in *nrg2*Δ::*kanMX4* strain. The CLA1-2 isogenic CLA1-3 (*rtg3*Δ) derivative was obtained by gene replacement. Two PCR-generated *kanMX* modules prepared from plasmid pFA6a (Supplementary Table S1) and four pairs of deoxyoligonucleotides: R1 - R2 (Supplementary Table S2a), were used to generate the corresponding *RTG3* module, following a previously described method (*Longtine et al., 1998*). Strain CLA1-2 was transformed with the 2612 bp PCR product containing the *kanMX4* cassette and *RTG3* upstream or downstream nucleotide sequences amplified from the genomic DNA of the CLA1-2 strain (Supplementary Table S2a).

**Table 1.**
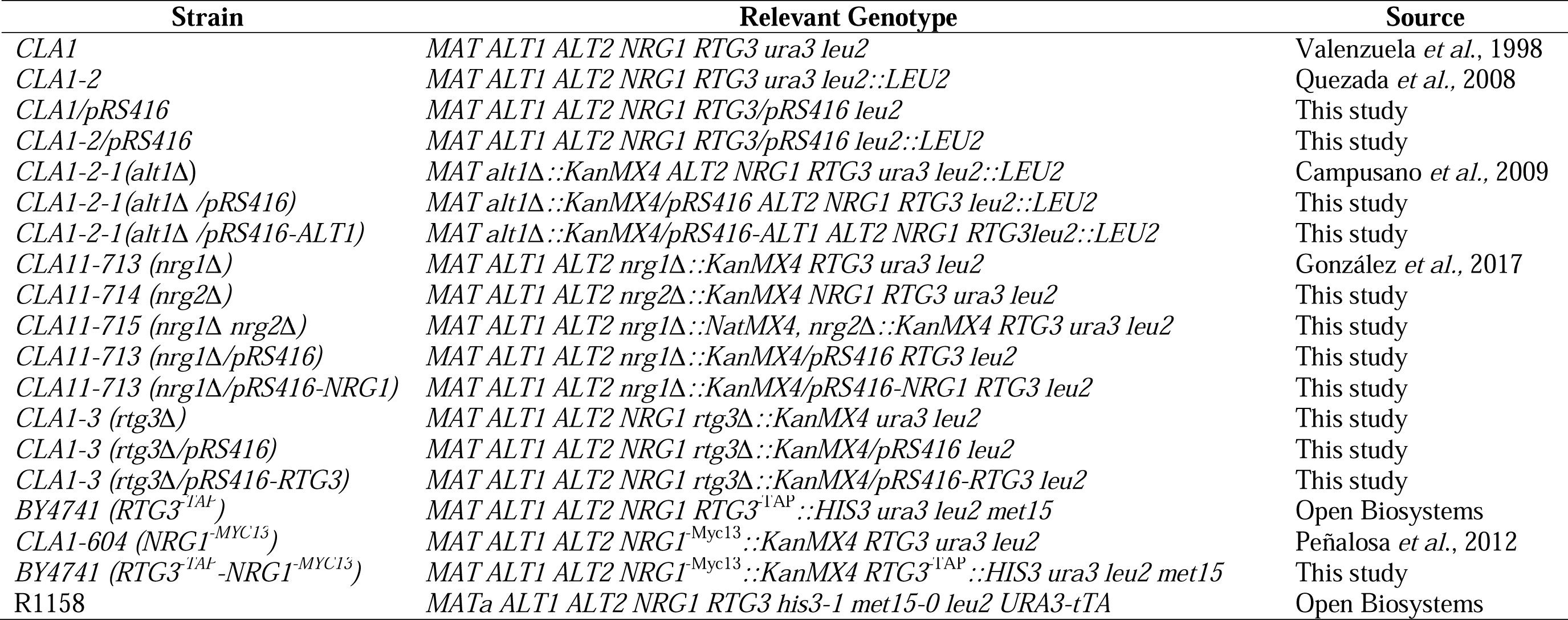
Strains used in the present work.

| Strain | Relevant Genotype | Source |
| --- | --- | --- |
| *CLA1* | *MAT ALT1 ALT2 NRG1 RTG3 ura3 leu2* | Valenzuela *et al.*, 1998 |
| *CLA1-2* | *MAT ALT1 ALT2 NRG1 RTG3 ura3 leu2::LEU2* | Quezada *et al.*, 2008 |
| *CLA1/pRS416* | *MAT ALT1 ALT2 NRG1 RTG3/pRS416 leu2* | This study |
| *CLA1-2/pRS416* | *MAT ALT1 ALT2 NRG1 RTG3/pRS416 leu2::LEU2* | This study |
| *CLA1-2-1(alt1Δ)* | *MAT alt1Δ::KanMX4 ALT2 NRG1 RTG3 ura3 leu2::LEU2* | Campusano *et al.*, 2009 |
| *CLA1-2-1(alt1Δ /pRS416)* | *MAT alt1Δ::KanMX4/pRS416 ALT2 NRG1 RTG3 leu2::LEU2* | This study |
| *CLA1-2-1(alt1Δ /pRS416-ALT1)* | *MAT alt1Δ::KanMX4/pRS416-ALT1 ALT2 NRG1 RTG3 leu2::LEU2* | This study |
| *CLA11-713 (nrg1Δ)* | *MAT ALT1 ALT2 nrg1Δ::KanMX4 RTG3 ura3 leu2* | González *et al.*, 2017 |
| *CLA11-714 (nrg2Δ)* | *MAT ALT1 ALT2 nrg2Δ::KanMX4 NRG1 RTG3 ura3 leu2* | This study |
| *CLA11-715 (nrg1Δ nrg2Δ)* | *MAT ALT1 ALT2 nrg1Δ::NatMX4, nrg2Δ::KanMX4 RTG3 ura3 leu2* | This study |
| *CLA11-713 (nrg1Δ/pRS416)* | *MAT ALT1 ALT2 nrg1Δ::KanMX4/pRS416 RTG3 leu2* | This study |
| *CLA11-713 (nrg1Δ/pRS416-NRG1)* | *MAT ALT1 ALT2 nrg1Δ::KanMX4/pRS416-NRG1 RTG3 leu2* | This study |
| *CLA1-3 (rtg3Δ)* | *MAT ALT1 ALT2 NRG1 rtg3Δ::KanMX4 ura3 leu2* | This study |
| *CLA1-3 (rtg3Δ/pRS416)* | *MAT ALT1 ALT2 NRG1 rtg3Δ::KanMX4/pRS416 leu2* | This study |
| *CLA1-3 (rtg3Δ/pRS416-RTG3)* | *MAT ALT1 ALT2 NRG1 rtg3Δ::KanMX4/pRS416-RTG3 leu2* | This study |
| *BY4741 (RTG3<sup>-TAP</sup>)* | *MAT ALT1 ALT2 NRG1 RTG3<sup>-TAP</sup>::HIS3 ura3 leu2 met15* | Open Biosystems |
| *CLA1-604 (NRG1<sup>-MYC13</sup>)* | *MAT ALT1 ALT2 NRG1<sup>-Myc13</sup>::KanMX4 RTG3 ura3 leu2* | Peñalosa *et al.*, 2012 |
| *BY4741 (RTG3<sup>-TAP</sup>-NRG1<sup>-MYC13</sup>)* | *MAT ALT1 ALT2 NRG1<sup>-Myc13</sup>::KanMX4 RTG3<sup>-TAP</sup>::HIS3 ura3 leu2 met15* | This study |
| R1158 | *MATa ALT1 ALT2 NRG1 RTG3 his3-1 met15-0 leu2 URA3-tTA* | Open Biosystems |

### Construction of Myc13-tagged strains

An *RTG3*-^TAP^ mutant was obtained from the BY4741 TAP-tagged *Saccharomyces* strain collection (BY4741 *ura3 leu2 his3 met5 RTG3-TAP*::*HIS3*). A BY44741 *RTG3-*^TAP^ *NRG1-*^Myc13^ derivative was obtained as previously described (*Longtine et al., 1998*). A pair of deoxyoligonucleotides: T1 and T2 (Supplementary Table S2b), was designed based on the *NRG1* coding sequence and that of the pFA6a-13Myc-*kanMX6* multiple cloning site (*Goldstein and McCusker 1999*), generating a PCR-13Myc*-kanMX6* module of 2300 bp used to transform the BY4741 *RTG3*-^TAP^ tagged strain, generating mutant strain *RTG3*-^TAP^ *NRG1*-^Myc13^ double tagged derivative. Deoxyoligonucleotides T3 and T4 (Supplementary Table S2b), were used to verify the construction *RTG3*^-TAP^ *NRG1*^-Myc13^. These primers generated a module of 2673 bp (233 bp of *NRG1* + 2300 bp of 13Myc*-kanMX6* +137 bp of 3’UTR of *NRG1*). Construction of the *NRG1*-^Myc13^ single strain mutant has been previously described (*Peñalosa et al., 2012*).

### Growth conditions

Strains were routinely grown on minimal medium (MM) containing salts, trace elements, and vitamins according to the formula for yeast nitrogen base (Difco). Glucose (2% w/v) or ethanol (2% v/v) were used as carbon sources. 7 mM GABA, 8 mM proline, 7mM alanine, or 40 mM ammonium sulfate were used as nitrogen sources. Uracil (20 mg/L), leucine or glutamate were added as auxotrophic requirements when needed. Cells were incubated at 30°C with shaking (250 rpm).

### *NRG1* and *RTG3* cloning

*ALT1* cloning has been previously described (*Rojas-Ortega et al., 2018*). For *NRG1* cloning, two variants of the *NRG1* containing module were created, the AB and the CD constructs (deoxyoligonucleotides are listed in Supplementary Table S2c). The AB region consists of a 2492 bp PCR product (deoxyoligonucleotides CN1 and CN2) harboring -1512 nucleotides from the *NRG1* start codon and +284 nucleotides from the *NRG1* stop codon. The CD region consists of the 2902 bp PCR product (oligonucleotides CN3 and CN4) harboring -1867 nucleotides from the *NRG1* start codon and +339 nucleotides from the *NRG1* stop codon. Recognition sites for endonuclease restriction enzymes *BamHI* and *XhoI*, were respectively added to the forward and reverse deoxyoligonucleotides, *NRG1* PCR products and pRS416 (*CEN6 ARS4 URA3*) plasmid (Supplementary Table S1), were *BamHI* - *XhoI* digested and ligated after gel purification. Ligations were transformed into the *DH5*α *Escherichia coli* strain. Plasmids were purified from *E. coli* extracts, and after correct cloning was verified by sequencing, plasmids were transformed into *S. cerevisiae* strains CLA1 *NRG1* and CLA11-713 *nrg1*Δ. Transformants were selected for uracil prototrophy on minimal medium (MM).

For *RTG3* cloning, a 2769 bp PCR product using deoxyoligonucleotides CR1and CR2 (Supplementary Table S2c) was generated, harboring -1089 nucleotides from the start codon and +219 nucleotides from the *RTG3* stop codon. Recognition sites for endonuclease restriction enzymes *BamHI* - *XhoI* were added to the forward and reverse deoxyoligonucleotides, respectively. *RTG3* PCR product and pRS416 (*CEN6 ARS4 URA3*) plasmid (Supplementary Table S2), were *BamHI* - *XhoI* digested and ligated after gel purification. Ligations were transformed into the *DH5a Escherichia coli* strain. Plasmids were purified, and correct cloning was verified by sequencing. Construct cloned in pRS416 was transformed into *S. cerevisiae* strains CLA1 *RTG3* and CLA1-3 *rtg3*Δ. Transformants were selected for uracil and glutamic acid prototrophy on minimal medium (MM).

### Northern blot analysis

Northern blot analysis was carried out as previously described (*González et al., 2017*). Total yeast RNA was prepared from 200 mL cultures grown to indicated OD_600nm_. Probes to monitor the expression of *ALT1, ALT2, CIT2*, *HXT2,* and *ACT1* were prepared by PCR from CLA1-2 genomic DNA using primers N1 to N10 (Supplementary Table S2d) and radioactively labeled by P^32^ with Random Primer Labeling Kit (Agilent, 300385). Blots were scanned using Image Quant 5.2 software (Molecular Dynamics).

### qPCR Differential Transcript Expression Analysis

*S. cerevisiae* strain was grown on MM Glucose (2% w/v) as carbon source and supplemented with GABA 7 mM or GABA + Ala 7 mM as nitrogen sources to an OD_600nm_ = 0.3. Cells were then harvested by centrifugation at 3,000 rpm × 5 min and washed twice with distilled H_2_O. After that, cells were transferred to a mortar, frozen with liquid nitrogen, and mechanically disrupted with the aid of a pestle. Then, total RNA was extracted with TRIzol (Invitrogen) following the TDS procedure. RNA quality and concentration were determined using a NanoDrop 2000 (NanoDrop Technologies). RNA was treated with RQ1 RNAse-Free DNAse (Promega) to degrade DNA from the samples; afterwards, cDNA was obtained using the RevertAid H Minus First Strand cDNA Synthesis Kit following manufacturer’s instructions (Thermo Scientific). cDNA samples were adjusted to a concentration of 30 ng/µL. PCR reactions were prepared with KAPA SYBR Fast kit (Roche) and run on the Corbett Research Rotor-Gene 6000 (Qiagen). Gene expression was obtained for the *ALT2*, *COX2*, *COX3*, *COB1*, *ATP9*, *ATP6*, *COX8*, and *COX6* genes, using deoxyoligonucleotides P1 to P18 (Supplementary Table S2e). Data were analyzed using the 2^^-ΔΔCT^ method in which RDN18S (18S) constitutive gene was used as housekeeping control (*Jozefczuk and Adjaye 2011*). The statistical analysis was made on GraphPad Prism 9.0 (GraphPad Software Inc). All experiments were repeated three times, and the results were expressed as mean + SEM.

### Nucleosome Scanning Assay

A nucleosome scanning assay (NuSA) was carried out as previously described (*Infante et al., 2012*). 100 mL cultures containing glucose (2% w/v) as carbon source and supplemented with GABA 7 mM or GABA + Ala 7 mM as nitrogen sources were grown at OD_600nm_ = 0.3. To determine nucleosome position on *ALT2* promoter, samples were treated as previously described (*González et al., 2017*). qPCR analysis was performed using a Corbett Life Science Rotor Gene 6000 and SYBR Green as dye (2X KAPA SYBR FAST, Invitrogen). Real-time PCR was carried out as follows: 94°C for 5 min (1 cycle), 94°C for 15s, 58°C for 20s, and 72°C for 20s (35 cycles). PCR deoxyoligonucleotides (B1 to B23) for *ALT2* analysis are described in supplementary Table S2g. The relative protection was calculated as a ratio to the control of *VCX1* that was amplified using P19 and P20 oligonucleotides (Supplementary Table S2e).

### Immunoprecipitation of Rtg3-^TAP^ and Nrg1-^Myc13^

The method was carried out using a modified version from *Gerace and Moazed (2014)*. *S. cerevisiae* strains were grown on minimum medium (MM) with glucose (2% w/v) as carbon source supplemented with γ-aminobutyric acid (GABA) 7 mM, GABA + Alanine 7 mM or GABA + Proline 8 mM as nitrogen sources at OD_600nm_ = 0.5, cells were harvested by centrifuging at 3000 rpm × 5 min and washed with TBS 1X. Later, microtubes (2mL) with the pellets were frozen, submerging them in liquid nitrogen for 15 s. 500 µL of lysis buffer (HEPES 50 mM, NaOAc 200 mM, EDTA 0.1 mM, EGTA 0.1 mM, MgOAc 5 mM, Glycerol 5%, NP-40 0.25%, DTT 3 mM, PMSF 1 mM, Protease inhibitor cocktail) and 400 µL of 0.5 mm glass bead were added and vortexed for 4 min twice with a pause of 2 min on ice. The microtube was centrifugated at 14,000 rpm for 5 min at 4°C; the total protein lysate was transferred to a new 2 mL microtube. Protein concentration was measured with the Bradford method. The normalized protein concentration (2000 µg or 4000 µg) was added to a microtube containing 60 µL of Pierce Protein A/G agarose beads (Thermo Scientific) and 10 µL of TAP Antibody CAB1001 (Invitrogen), 10 µL of Myc13 Antibody 9E11 (Santa Cruz Biotechnology) or 10 µL of HA antibody sc-7392 (Santa Cruz Biotechnology). Samples were incubated on mild agitation for 2 hrs at 4°C. After, samples were washed 5 times with 1 mL of wash buffer (HEPES 50 mM, NaOAc 200 mM, EDTA 0.1 mM, EGTA 0.1 mM, MgOAc 5 mM, Glycerol 5%, NP-40 0.25%, DTT 3 mM, PMSF 1 mM) centrifuging at 2,000 rpm for 1 min at 4°C. Subsequently, all supernatant was removed, and 50 µL of Loader buffer (Tris base 50 mM pH 6.8, SDS 2%, Glycerol 10%, DTT 100 mM, PMSF 1 mM, bromophenol blue 0.001%) was added, and incubated at 65°C for 10 min. 40 µl were run on 10% SDS-PAGE for Western blotting analysis using anti-TAP antibodies (Thermo Scientific).

### Quantitative chromatin immunoprecipitation

Formaldehyde cross-linking and immunoprecipitations were carried out adapting a previously described procedure (*Hernández et al., 2011*). Yeast cells (200 mL of OD_600nm_ = 0.3) were cross-linked with 1% formaldehyde for 20 min at room temperature. Afterward, 125 mM glycine was added and incubated for 5 min. Cells were then harvested and washed with PBS buffer. Pelleted cells were suspended in lysis buffer (140 mM NaCl, 1 mM EDTA, 50 mM HEPES/KOH, 1% Triton X-100, 0.1% sodium deoxycholate) with a protease inhibitor cocktail (Complete Mini, Roche). Cells were lysed with glass beads and collected by centrifugation. Extracts were sonicated with a Diagenode Bioruptor to produce chromatin fragments with an average size of 300 bp. Immunoprecipitation reactions were carried out with 1 mg anti-c-Myc antibody (9E11, Santa Cruz Biotechnology) and protein A beads for 3 hrs, washed, suspended in TE buffer/1% SDS, and incubated overnight at 65°C to reverse the formaldehyde cross-linking. Immunoprecipitates were then incubated with proteinase K (Roche), followed by phenol/chloroform/isoamyl alcohol extraction, precipitation, and suspension in 30 µl TE buffer. Dilutions of input DNA (1:100) and immunoprecipitated DNA (1:2) were analyzed by qPCR. Real-time PCR-based DNA amplification was performed using specific primers that were initially screened for dimer absence or cross-hybridization. Only primer pairs (C1 to C18) with similar amplification efficiencies were used (Supplementary Table S2f). Quantitative chromatin immunoprecipitation (qChIP) analysis was performed using a Corbett Life Science Rotor Gene 6000 machine. The fold difference between immunoprecipitated material (IP) and total input sample for each qPCR-amplified region was calculated following the formula IP/input = (2InputCt 2 IPCt) (*Litt et al., 2001*). The results represent the mean values and SE of at least three independent, crosslinked samples, each being immunoprecipitated twice with the antibody.

### Ethanol measurement

For ethanol, determination cells were grown to indicate OD_600nm_ in glucose and ammonium sulfate as nitrogen source. Aliquots of 1 mL were withdrawn for supernatant recovery by centrifugation at 14,000 rpm. Ethanol quantification was determined in the supernatant following a previously reported protocol (*Calahorra et al., 2012*) using the reaction buffer (bicine-KOH 20 mM, pH=9.0 and 1 U/mL alcohol dehydrogenase, Sigma Aldrich A7011). The assay was carried out at 340 nm, and 25 °C in sample concentration was used (*Márquez et al., 2021*).

### Fluorescent microscopy

Cells were stained with Green MitoTracker FM 9074 (Molecular Probes) according to manufacturer specifications. Confocal images were obtained using a FluoView FV1000 laser confocal system (Olympus) attached/interfaced to an Olympus IX81 inverted light microscope with a 60x oil-immersion objective (UPLASAPO 60x O NA: 1.35), zoomx20.0 and 3.5 μm of confocal aperture. The excitation and emission settings were as follows: MitoTracker excitation 543 nm; emission 598 nm, BF 555 nm range 100 nm. The subsequent image processing was carried out with Olympus Fluo View FV1000 (version324 1.7) software.

### Quantification of mtDNA/nDNA ratio

The amount of mtDNA relative to nDNA was obtained to quantify the mtDNA. We followed the method described by *Márquez et al., 2021*, estimating mtDNA/nDNA ratio by qPCR using different genes of mtDNA: *COX2*, *COX3,* and *ATP6*, and as a nuclear-encoded gene, we selected *COX6*. Total DNA from the indicated strains was extracted twice by phenol/chloroform with 20 µl of NaCl 5 M. Samples were incubated with 20 µg RNase A for 1 hr at 37°C. DNA precipitation was carried out with an equal volume of ethanol for 30 min at -20°C and resuspended in nuclease-free water. DNA was quantified by NanoDrop, ThermoScientific and samples were diluted at 50 ng/µL for qPCR assay. qPCR was performed using SYBR Green as dye (2X KAPA SYBR FAST, Invitrogen). Conditions for qPCR were: 94°C for 5 min (1 cycle), 94°C for 15 s, 59°C for 30 s, and 72°C for 20 s (30 cycles). We obtained the Ct (cycle threshold) for each sample, and following the ΔΔCt method, we calculated the mtDNA/nDNA ratio as described previously (*Quiros et al., 2017*).

### Phylogeny of yeast proteins

The PhylomeDB (http://phylomedb.org/) database was used (*Fuentes et al., 2022*) to analyze Nrg1 and Nrg2 phylogeny.

### Structural modeling and visualization

Extant structural models were downloaded from the alpha-fold database (https://alphafold.ebi.ac.uk/) (*Varadi et al., 2022*). The model of the chimeric complex was obtained through AphaFold2.ipynb (https://colab.research.google.com/github/sokrypton/ColabFold/blob/main/AlphaFold 2.ipynb#scrollTo=ADDuaolKmjGW) (*Mirdita et al., 2022*).

Contacts between chains were determined using the Mapiya Contact Map Server (https://mapiya.lcbio.pl/; *Badaczewska-Dawid et al., 2022*). Structural models were visualized and figures drawn with ChimeraX-1.5 for Nrg1-Nrg2 chain comparison and with VMD 1.9.4 for visualization of the Nrg1-Rtg3 complex and inter-chain contacts (*Humphrey et al., 1996; Goddard et al., 2018; Pettersen et al., 2021)*. http://www.ks.uiuc.edu/Research/vmd/

## Acknowledgements

We are grateful Manuel L Robert for motivating discussions which influenced our work, to Félix Recillas Targa for his support and his critical review of the manuscript and to Jesús Aguirre for fruitful discussions, along the development of this work. To Ruth Rincón-Heredia, Head of the Imaging Unit, Institute of Cell Physiology, and Rosario Ortiz-Hernández from the Cell Biology Department of the Faculty of Sciences UNAM for microscopy analysis assistance, to Laura Ongay, Guadalupe Códiz and Minerva Mora from the Molecular Biology Unit, IFC, UNAM and to Sheila Cruz Cruz for skillful technical assistance. We are also indebted to Rocio Romualdo Martínez for helpful secretarial assistance.

## Additional information

### Author contributions

LR-R, JG, CS, and AG designed research, analyzed data and wrote the manuscript. CC-B, JG, ER, HH, BA, NT-R, DM, NS, NG-H performed research. All authors contributed to the article and approved the submitted version.

### Author ORCIDS

Carlos Campero-Basaldua 0009-0005-7296-2705; James González 0000-0002-7329-4025; Janeth García-Rodriguez 0009-0008-6647-6235; Edgar Ramírez 0000-0002-5496-0846; Hugo Hernandez 0000-0002-4315-6799; Beatriz Aguirre 0000-0002-3288-7967; Nayeli Torres-Ramírez 0000-0002-7515-9740; Dariel Márquez 0000- 0001-9988-3912; Norma S Sánchez 0000-0002-9954-5451; Nicolás Gómez-Hernández 0000-0002-8773-5258; Lina Riego-Ruiz 0000-0002-2883-8871; Claudio Scazzocchio 0000-0001-8800-2534; Alicia González 0000-0002-1276-5719.

