## Supplementary Figures for "Neo-functionalization in *Saccharomyces cerevisiae*: A Novel Nrg1-Rtg3 chimeric transcriptional modulator is essential to maintain mitochondrial DNA integrity"

**Supplementary Figure 1.**

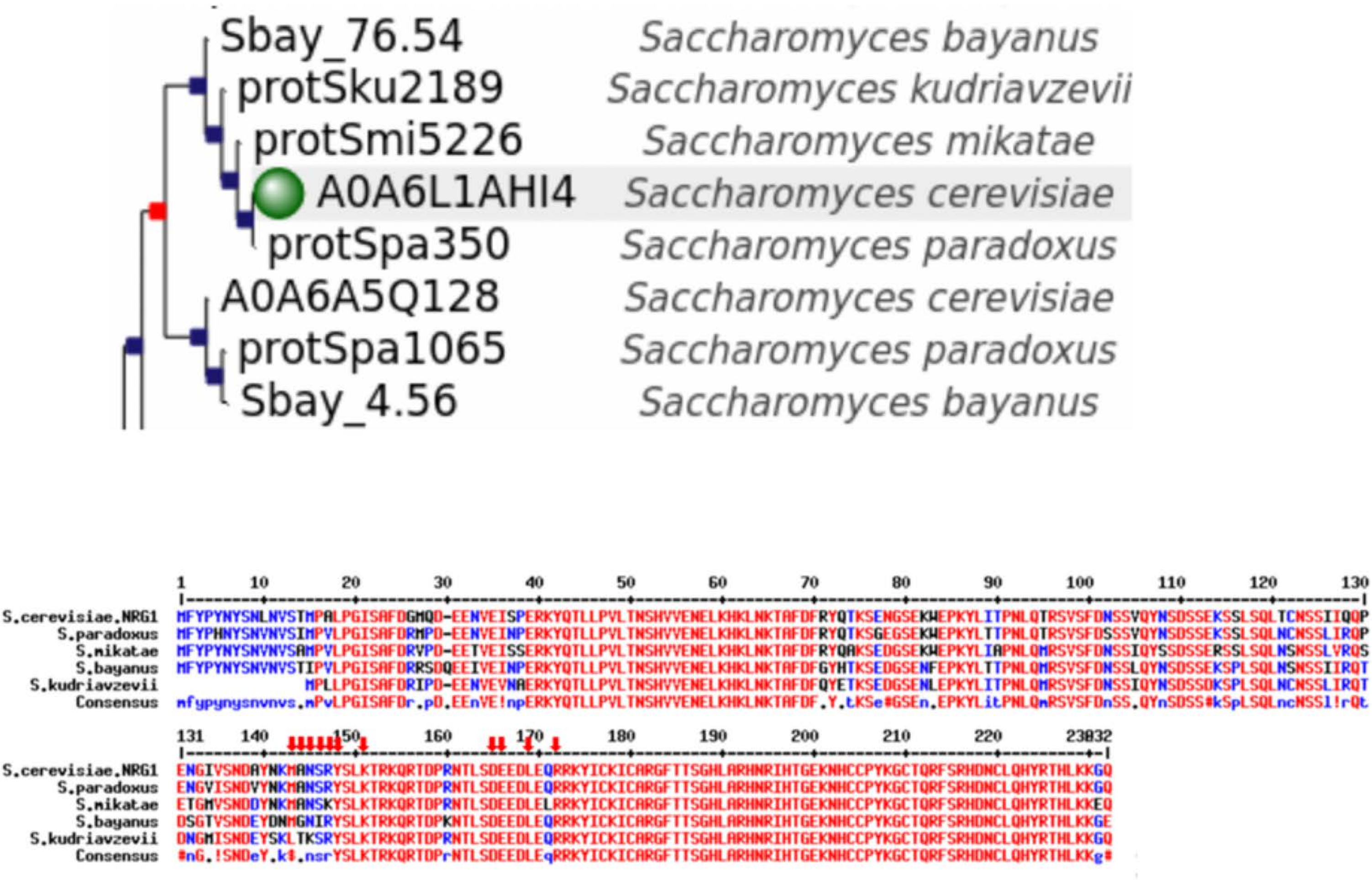

### Supplementary Figure 2.

```
           1      10        20        30        40        50        60        70        80        90        100       110       120       130
           |--------+---------+---------+---------+---------+---------+---------+---------+---------+---------+---------+---------+---------|
NRG1              MFYPYNYSNLNVSTMPALPGISAFDGMQDEENVEISPERKYQTLLPVLTNSHVVENELKHKLNKTAFDFRYQTKSENGSEKWEPKYLITPNLQTRSVSFDNSSVQYNSDSSEKSSLSQLTCNS
NRG2       MSIGYKDNLMSTILAKDRKCEFPINFECSPSQITLMPEMFSFNNERKYQTLIPLMKTSHLIDDDLKDKLNKCAFDF-FSGKQANRTS---------------------DGTISRLTASGKTSPILPLQ-NI
Consensus  .......nlmptnlandrkcemPanfecSaf#gmqdeEnfeinnERKYQTLiPl$knSHl!###LKdKLNKcAFDF.%qgKqaNrse.....................#gs!qrnsaSgekSpilqLq.Ni

           131    140       150       160       170       180       190       200       210       220       230       240      248
           |--------+---------+---------+---------+---------+---------+---------+---------+---------+---------+---------+--------|
NRG1       SIIQQPENGIVSNDAYNKMANSRYSLKTRKQRTDPRNTLSDEEDLEQRRKYICKICARGFTTSGHLARHNRIHTGEKNHCCPYKGCTQRFSRHDNCLQHYRTHLKKGQ
NRG2       NIVKAENTGNGKSDPYSSIKISKPTKTVIKLKSTKTNTAG------QRTRHFCKICSTGFTTSGHLSRHNRIHTGEKNHICPHEGCGQRFSRHDNCNQHYRTHANKKKRNWKRREASS
Consensus  nI!qae#nGngknDaYnkianSrpslktrKqrsdkrNTag......QRrrhiCKICarGFTTSGHLaRHNRIHTGEKNHcCPheGCgQRFSRHDNCnQHYRTHanKgq..........
```

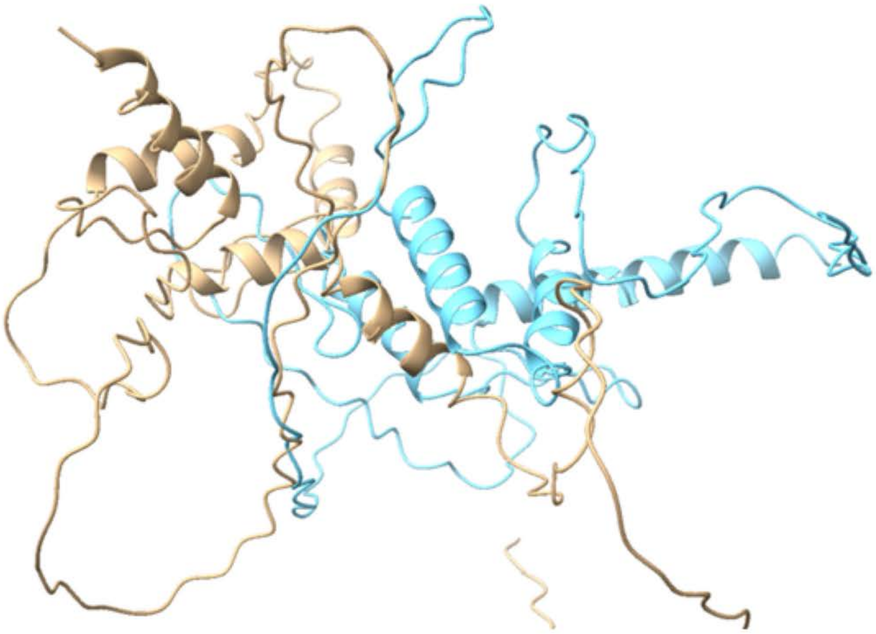

Gold: NRG1
Cyan: NRG2

#### Supplementary Figure 3.

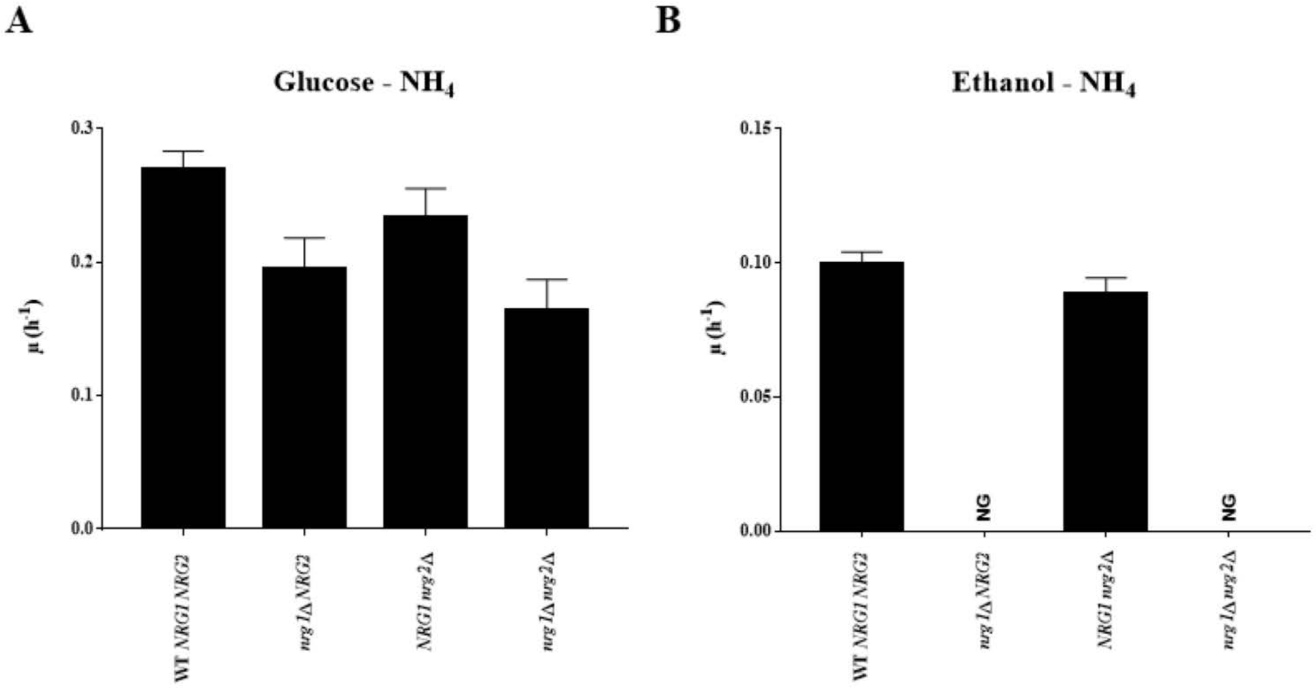

**SUPPLEMENTARY FIGURE 1. Comparative analysis of *S. cerevisiae* orthologues**. Top panel, clade comprising the orthologues of the *S. cerevisiae* Nrg1 protein (marked with a green sphere extracted from PhylomeDB (http://phylomedb.org/), duplication event shown as a red dot. Bottom panel, alignment of the orthologues comprised in the clade, with putative residues of contact of Nrg1 with Rtg3 indicated by red arrows. See Figure 9 of the main text for details concerning contact modelling and visualization.

**SUPPLEMENTARY FIGURE 2. Nrg1 and Nrg2 Comparison. This figure highlights the differences between the two paralogues.** Top panel sequence comparison. The two patches involved in contacts of Nrg1 with Rgt3 are highlighted with red rectangles. Bottom, Alpha-fold models of Nrg1 and Nrg2 obtained from the Alpha-fold data base (https://alphafold.ebi.ac.uk/) and compared with Matchmaker in ChimeraX-1.5 (*Pettersen et al.*, 2021; *Goddard et al.*, 2018). Note that only one α-helix is superimposed.

**SUPPLEMENTARY FIGURE 3. *nrg2*Δ mutants show wild type growth on ethanol, while *nrg1*Δ *nrg2*Δ double mutants do not grow on ethanol.** (A) Specific growth rate on MM glucose (2%)-ammonium (40 mM), (B) specific growth rate on MM ethanol (2%)-ammonium (40mM). Results of three independent experiments are presented.
