## Supplementary Tables for "Neo-functionalization in *Saccharomyces cerevisiae*: A Novel Nrg1-Rtg3 chimeric transcriptional modulator is essential to maintain mitochondrial DNA integrity"

**Table S1. Plasmids used in this study.**

| Plasmids | Source |
|---|---|
| pFA6a-KanMX4 | Longtine MS *et al.* 1998 |
| p4339 | Goldstein and McCusker 1999 |
| pFA6a-Myc13-kanMX6 | Longtine MS *et al.* 1998 |
| pRS416 | Sikorski RS and Hieter P *et al.* 1989 |

**Table S2. Deoxyoligonucleotides used in the present work.**

**a)** Deoxyoligonucleotides used to construct null mutants.

| Name | Sequence (5' to 3') | Application |
|---|---|---|
| R1 | ATTTTTTGTCAGGCGAACCTACTTCTTAAATAAGTGAAGA**CGTACGCTGCAGGTCGACGGA** | Fw from -1 to -40 of *RTG3* and for pFA6A-KanMx4 (sequence in bold) used to generate *rtg3*Δ mutant |
| R2 | TACCTGACCTTTTTCAAATTTAATTTTTTCCCGCTAATAA**ATCGATGAATTCGAGCTCGT** | Rv from +2471 to +2510 of *RTG3* and for pFA6A-KanMx4 (sequence in bold) used to generate *rtg3*Δ mutant |
| R3 | ATGTTTTACCCATATAACTATAGTAACCTCAATGTTTCTACTATGCCCGCAC**CGTACGCTGCAGGTCGAC** | Fw from +1 to +51 of *NRG1* and for pFA6A-KanMx4 (sequence in bold) used to generate *nrg1*Δ mutant |
| R4 | TTATTGTCCCTTTTTCAAATGTGTTCTATAGTGTTGCAAGCAATTATCATG**ATCGATGAATTCGAGCTCG** | Rv from +635 to +693 of *NRG1* and for pFA6A-KanMx4 (sequence in bold) used to generate *nrg1*Δ mutant |
| R5 | GACAGCTCAAATGAATTTCCGGCACCAAGTCATATGAGCAC**CGTACGCTGCAGGTCGAC** | Fw from -50 to -91 of *NRG2* and for pFA6A-KanMx4 (sequence in bold) used to generate *nrg2*Δ mutant |
| R6 | GAACCGACTATCGTTCCTGCTATTGTTCTTGGTGAGTATTAC**ATCGATGAATTCGAGCTCG** | Rv from +751 to +792 of *NRG2* and for pFA6A-KanMx4 (sequence in bold) used to generate *nrg2*Δ mutant |

***b)*** Deoxyoligonucleotides used to tag with the Myc13 epitope the Strain *RTG3*$^{-TAP}$- *NRG1*$^{-Myc13}$.

| Name | Sequence (5' to 3') | Application |
|---|---|---|
| T1 | GACATGATAATTGCTTGCAACACTATAGAACACATTTGAAAAAGGGACA**ACGGATCCCCGGGTTAATTAA** | Fw from +643 to +693 of *NRG1* and for pFA6A-KanMx4-13Myc (sequence in bold) used to generate *NRG1*$^{-Myc13}$ tagged |
| T2 | GTAAAGTGCGGAATAGTAGTACTGCTAATGAGAAAAACACGGGTATACCGTCAA**GAATTCGAGCTCGTTTAAAC** | Rv from +697 to +750 of *NRG1* and for pFA6A-KanMx4-13Myc (sequence in bold) used to generate *NRG1*$^{-Myc13}$ tagged |
| T3 | AAAGAACAGATCCAAGAAATACG | Fw from +460 to +483 of *NRG1* to verify Myc-$^{13}$-*kanMX4* tagging on *NRG1* |
| T4 | TCTGAAAGTGATTTATGATGCCA | Rv from +810 to +833 of *NRG1* to verify Myc-$^{13}$-*kanMX4* tagging on *NRG1* |

**c)** Deoxyoligonucleotides used to *NRG1* or *RTG3* cloning.

| **Name** | **Sequence (5' to 3')** | **Application** |
|---|---|---|
| CN1 | ATC**GGATCC**AAGCTGCGCCTATATGCCTT | Fw from -1512 to -1492 of *NRG1* and BamHI restriction site (sequence in bold) used to clone into the pRS416 |
| CN2 | TGG**CTCGAG**AGCCTTGTTGCAGCACTACT | Rv from +957 to +977 of *NRG1* and XhoI restriction site (sequence in bold) used to clone into the pRS416 |
| CN3 | GTTT**GGATCC**ACAATCGTGCCATTTAACCGG | Fw from -1867 to -1846 of *NRG1* and BamHI restriction site (sequence in bold) used to clone into the pRS416 |
| CN4 | ACAG**CTCGAG**AGGAGGTAGTCACAGTCTCGT | Rv from +1012 to +1032 of *NRG1* and XhoI restriction site (sequence in bold) used to clone into the pRS416 |
| CR1 | TTC**GGATCC**AGCACATGTAATAGTCAGC | Fw from -1089 to -1069 of *RTG3* and BamHI restriction site (sequence in bold) used to clone into the pRS416 |
| CR2 | TCT**CTCGAG**CTATCTCTTCCACTCTTTTC | Rv from +1660 to +1680 of *RTG3* and XhoI restriction site (sequence in bold) used to clone into the pRS416 |

**d)** Deoxyoligonucleotides used to Northern blot assays.

| Name | Sequence (5' to 3') | Application |
|---|---|---|
| N1 | AGACCCGTCCTACAGAGACATAGC | Fw from +172 to +195 of *ALT1* used for Northern probe |
| N2 | GCGAGCTTCTTGAACTGCCTTGAA | Rv from +1558 to +1581 of *ALT1* used for Northern probe |
| N3 | GACACACCAACAGGATTTGAAAGG | Fw from +8 to +32 of *ALT2* used for Northern probe |
| N4 | GCGTTCGCTTTCACAAAGAGCTT | Rv from +1306 to +1329 of *ALT2* used for Northern probe |
| N5 | AGCGAAATCTACCCCATCCATGCTC | Fw from +91 to +115 of *CIT2* used for Northern probe |
| N6 | CCTTTCAATGGAAGCACCGATGGC | Rv from +1297 to +1320 of *CIT2* used for Northern probe |
| N7 | CTGAACTCCCAGCAAAGCCAATCG | Fw from +127 to +141 of *HXT2* used for Northern probe |
| N8 | CAAACAGCCCATGAAGACATACCCG | Rv from +1455 to +1469 of *HXT22* used for Northern probe |
| N9 | GTTTTGCCGGTGACGAC | Fw from +368 to +384 of *ACT1* used for Northern probe |
| N10 | CTTTCGGCAATACCTGGG | Rv from +1227 to +1244 of *ACT1* used for Northern probe |

**e)** Deoxyoligonucleotides used to qPCR assays.

| Name | Sequence (5' to 3') | Application |
|---|---|---|
| P1 | TCTCAAACAGGCCTGCTGAT | Fw from 562 to 581 of *ALT2* used for qPCR analysis |
| P2 | TCGTCGCTGTTTGTAGACCA | Rv from 689 to 670 of *ALT2* used for qPCR analysis |
| P3 | CATTCAGCTATGAGTCCTG | Fw from 330 to 348 of mitochondrial gene *COX3* used for qPCR analysis |
| P4 | GTTACAGTAGCACCAGAAGA | Rv from 457 to 438 of mitochondrial gene *COX3* used for qPCR analysis |
| P5 | GATGCTACTCCTGGTAGATT | Fw from 591 to 610 of mitochondrial gene *COX2* used for qPCR analysis |
| P6 | GTCCCACACAACTCAGAA | Rv from 679 to 662 of mitochondrial gene *COX2* used for qPCR analysis |
| P7 | CGCTAGAGCTATTTCATTAG | Fw from 530 to 549 of mitochondrial gene *ATP6* used for qPCR analysis |
| P8 | GGTACAAAACCGAATACTAA | Rv from 664 to 645 of mitochondrial gene *ATP6* used for qPCR analysis |
| P9 | TGGAGCAGGTATCTCAAC | Fw from 29 to 46 of mitochondrial gene *ATP9* used for qPCR analysis |
| P10 | CCATAGGGAATACTAGGTCT | Rv from 150 to 131 of mitochondrial gene *ATP9* used for qPCR analysis |
| P11 | GTAGGTAACGATATTGTATC | Fw from 468 to 488 of mitochondrial gene *COB1* used for qPCR analysis |
| P12 | TAGATGAACCATGAATATGT | Rv from 622 to 602 of mitochondrial gene *COB1* used for qPCR analysis |
| P13 | ACACTTCAAAGACGGTGTGT | Fw from 84 to 103 of *COX8* used for qPCR analysis |
| P14 | TCCAATAGCGAAGAACCCGA | Rv from 180 to 161 of *COX8* used for qPCR analysis |

| | | |
|---|---|---|
| P15 | GCCAGACCTGGTGCTTATCATGC | Fw from +51 to +77 of *COX6* used for qPCR analysis and qPCR in mtDNA/nDNA ratio analysis |
| P16 | GGGAACGCCCAATTCTTGTCTGAC | Rv from +391 to +414 of *COX6* used for qPCR analysis and qPCR in mtDNA/nDNA ratio analysis |
| P17 | ACTTCTTAGAGGGACTATCG | Fw from 1374 to 1394 of *RDN18S* used for qPCR analysis |
| P18 | CAAGATTACCAAGACCTCTC | Rv from 1501 to 1521 of *RDN18S* used for qPCR analysis |
| P19 | TGCGTGTGCATCCCTACTGA | Fw from +504 to +523 of *VCX1* used to normalize DNA protection in NuSA experiments |
| P20 | AAGTGGTCTTCCTTGCCATGA | Rv from +552 to +572 of *VCX1* used to normalize DNA protection in NuSA experiments |

**f)** Deoxyoligonucleotides used for ChIP  of *ALT2* assay.

| Name | Sequence (5' to 3') | Application |
|---|---|---|
| C1 | TTCAAAGAAGTTGCCGCCGT | Fw from -821 to -801 of *ALT2* used for Nrg1 binding ChIP assay |
| C2 | TGTATGTGAATGAGCAGTGGAT | Rv from -732 to -710 of *ALT2* used for Nrg1 binding ChIP assay |
| C3 | ATCCACTGCTCATTCACATACA | Fw from -731 to -709 of *ALT2* used for Nrg1 binding ChIP assay |
| C4 | TGGGTGAGGCCGCAGATT | Rv from -629 to -611 of *ALT2* used for Nrg1 binding ChIP assay |
| C5 | GCTTCACCCTTGATTGGCAA | Fw from -568 to -548 of *ALT2* used for Nrg1 binding ChIP assay |
| C6 | AAGACGTGTGGAGGACATCA | Rv from -631 to -611 of *ALT2* used for Nrg1 binding ChIP assay |
| C7 | TACTTCGTCATAATTACTTCATTA | Fw from -443 to -419 of *ALT2* used for Nrg1 binding ChIP assay |
| C8 | TTGCGAATGAGGCCAAACTT | Rv from -360 to -340 of *ALT2* used for Nrg1 binding ChIP assay |
| C9 | TCCTACTTGTTTGGCGATTTTATT | Fw from -302 to -278 of *ALT2* used for Nrg1 binding ChIP assay |
| C10 | AAGTGTATTGGTAATTTGAAGAG | Rv from -235 to -212 of *ALT2* used for Nrg1 binding ChIP assay |
| C11 | TCTCTAGAGGCATAAAAGATAAG | Fw from -91 to -68 of *ALT2* used for Nrg1 binding ChIP assay |
| C12 | TGTCATTGTCATGTGTTCTTACTT | Rv from -12 to +12 of *ALT2* used for Nrg1 binding ChIP assay |
| C13 | CCGTAGAAAAGAAAAAGAACCGGGG | Forward deoxynucleotide to *HXT2* gene used as positive control for Nrg1 binding for ChiPassays |
| C14 | GGGGAAAGCAAAGCCACGTGGA | Reverse deoxynucleotide to *HXT2* gene used as positive control for Nrg1 binding for ChiP assays |

| | | |
|---|---|---|
| C15 | AGCGCGCACATTCGACGCATTTAT | Forward deoxynucleotide to *CIT2* gene used as positive control for Rtg3 binding for ChiP assays |
| C16 | GTCTAGCTACGGAAAAGGTCACAC | Reverse deoxynucleotide to *CIT2* gene used as positive control for Rtg3 binding for ChiP assays |
| C17 | AAGCGTTCGGTTGGATGAC | Forward deoxynucleotide to *GRS1* gene used as negative control for ChiP assays |
| C18 | GGGGAGCAAACACTATTCACTT | Reverse deoxynucleotide to *GRS1* gene used as negative control for ChiP assays |

**g)** Deoxyoligonucleotides used for *ALT2* Nucleosome Scanning Assay (NuSA).

| Name | Sequence 5' to 3' | Middle point of PCR (promoter coordinate) | 5'/3' end | Amplicon size (bp) |
|---|---|---|---|---|
| B1 | Fw TTCAAAGAAGTTGCCGCCGT<br>Rv TGTATGTGAATGAGCAGTGGAT | -766 | -821<br>-710 | 111 |
| B2 | Fw CTGAAGAACTAGAGGACTTAGT<br>Rv CAGGAGCAACCGCTTTTAAC | -701 | -760<br>-641 | 119 |
| B3 | Fw ATCCACTGCTCATTCACATACA<br>Rv TGGGTGAGGCCGCAGATT | -671 | -731<br>-611 | 120 |
| B4 | Fw GTTAAAAGCGGTTGCTCCTG<br>Rv TTGCCAATCAAGGGTGAAGC | -604 | -660<br>-548 | 112 |
| B5 | Fw ACCGCCATTGTTCGTACGT<br>Rv GAAACAATAGTATCGTGATGTGA | -555 | -600<br>-510 | 90 |
| B6 | Fw GCTTCACCCTTGATTGGCAA<br>Rv AAGACGTGTGGAGGACATCA | -520 | -568<br>-472 | 96 |
| B7 | Fw TCACATCACGATACTATTGTTTC<br>Rv ATAATGAAGTAATTATGACGAAG | -476 | -533<br>-418 | 115 |
| B8 | Fw TGATGTCCTCCACACGTCTT<br>Rv TTCTTAGGTATAATGAAGTAATT | -451 | -492<br>-409 | 83 |
| B9 | Fw TACTTCGTCATAATTACTTCATTA<br>Rv TTGCGAATGAGGCCAAACTT | -392 | -443<br>-340 | 103 |
| B10 | Fw AATTACTTCATTATACCTAAGAA<br>Rv ACAAGCCGTTTCCATGGAGA | -374 | -432<br>-315 | 92 |
| B11 | Fw AAGTTTGGCCTCATTCGCAA<br>Rv AATAAAATCGCCAAACAAGTAGG | -319 | -360<br>-278 | 82 |
| B12 | Fw TCTCCATGGAAACGGCTTGT<br>Rv TATGGGACCATAAGAGTAGGA | -293 | -335<br>-250 | 85 |

| | | | | |
|---|---|---|---|---|
| B13 | Fw TCCTACTTGTTTGGCGATTTTATT<br>Rv AAGTGTATTGGTAATTTGAAGAG | -257 | -302<br>-212 | 90 |
| B14 | Fw TCCTACTCTTATGGTCCCATA<br>Rv TTCCCACACGAAGTAGTAAATTT | -225 | -271<br>-178 | 93 |
| B15 | Fw ATCTCTTCAAATTACCAATACACT<br>Rv AAAGGATCAAAAAGAAAAGTGA | -196 | -247<br>-145 | 102 |
| B16 | Fw AAATTTACTACTTCGTGTGGGAA<br>Rv TATTTACTGAATGAGAAAAGGAT | -155 | -201<br>-109 | 92 |
| B17 | Fw TGTTCACTTTTCTTTTTGATCCTTT<br>Rv TCTTATCTTTTATGCCTCTAGAGA | -119 | -170<br>-67 | 103 |
| B18 | Fw TCATCCTTTTCTCATTCAGTAAAT<br>Rv ACTTACAAAAACAACACTACTGT | -59 | -134<br>-26 | 108 |
| B19 | Fw TCTCTAGAGGCATAAAAGATAAG<br>Rv TGTCATTGTCATGTGTTCTTACTT | -52 | -91<br>+12 | 103 |
| B20 | Fw CACAGTAGTGTTGTTTTTGTAAGT<br>Rv TCCTTTGCGGTGAACACAC | +1 | -50<br>+50 | 100 |
| B21 | Fw AAGTAAGAACACATGACAATGAC<br>Rv TAATCTTGCCAGCGGGCTTA | +34 | -12<br>+79 | 91 |
| B22 | Fw GTGTGTTCACCGCAAAGGA<br>Rv AGCATATTCTGCCTTAGTGACA | +72 | +31<br>+113 | 82 |
| B23 | Fw TAAGCCCGCTGGCAAGATTA<br>Rv TTAGCTCGTCAGCTCTGGTT | +110 | +59<br>+160 | 101 |

**Supplementary Table 3. Amino acid molecular contacts and interactions forces between Nrg1 and Rtg3.**

| Nrg1 | Rtg3 | distance | possible_interaction_forces |
|---|---|---|---|
| MET:142 | SER:188 | 5.808 | electrostatic: dipole-dipole, hydrogen bond, |
| MET:142 | SER:189 | 7.679 | electrostatic: dipole-dipole, hydrogen bond, |
| MET:142 | SER:190 | 7.474 | electrostatic: dipole-dipole, hydrogen bond, |
| MET:142 | ASN:192 | 7.157 | electrostatic: dipole-dipole, hydrogen bond, |
| ALA:143 | SER:188 | 6.495 | Induction + dispersion |
| ASN:144 | LEU:187 | 6.062 | Induction + dispersion |
| ASN:144 | SER:188 | 3.205 | electrostatic: dipole-dipole, hydrogen bond, |
| ASN:144 | SER:189 | 5.754 | electrostatic: dipole-dipole, hydrogen bond, |
| ASN:144 | SER:190 | 6.506 | electrostatic: dipole-dipole, hydrogen bond, |
| SER:145 | SER:185 | 5.624 | electrostatic: dipole-dipole, hydrogen bond, |
| SER:145 | SER:186 | 5.802 | electrostatic: dipole-dipole, hydrogen bond, |
| SER:145 | LEU:187 | 3.695 | Induction + dispersion |
| SER:145 | SER:188 | 3.061 | electrostatic: dipole-dipole, hydrogen bond, |
| SER:145 | SER:189 | 6.882 | electrostatic: dipole-dipole, hydrogen bond, |
| ARG:146 | LEU:187 | 6.101 | Induction + dispersion |
| ARG:146 | SER:188 | 5.412 | electrostatic: dipole-dipole, hydrogen bond, |
| TYR:147 | LEU:183 | 7.254 | hydrophobic, |
| TYR:147 | SER:184 | 5.619 | electrostatic: dipole-dipole, hydrogen bond, dipole- π stacking |
| TYR:147 | SER:185 | 2.96 | electrostatic: dipole-dipole, hydrogen bond, dipole- π stacking |
| TYR:147 | SER:186 | 2.922 | electrostatic: dipole-dipole, hydrogen bond, dipole- π stacking |
| TYR:147 | LEU:187 | 4.509 | hydrophobic, |
| TYR:147 | SER:188 | 6.252 | electrostatic: dipole-dipole, hydrogen bond, dipole- π stacking |
| LYS:150 | ASP:182 | 4.816 | salt bridge, hydrogen bond, |
| LYS:150 | LEU:183 | 4.574 | hydrophobic, |
| LYS:150 | SER:184 | 7.419 | electrostatic: dipole-dipole, hydrogen bond, |
| ASP:164 | LYS:327 | 2.669 | salt bridge, hydrogen bond, |
| GLU:165 | LYS:327 | 7.573 | salt bridge, hydrogen bond, |
| LEU:168 | LYS:327 | 7.248 | hydrophobic, |
| ARG:171 | LYS:327 | 7.011 | Ionic repulsion CAUTION: possible repulsion |
